# *SRRM2* haploinsufficiency drives SynGAP-γ reduction, *AGAP3* mis-splicing, and oligodendrocyte deficits in a mouse model of schizophrenia

**DOI:** 10.1101/2024.10.10.617460

**Authors:** Sameer Aryal, Chuhan Geng, Margaret A. Yin, Min Jee Kwon, Zohreh Farsi, Nathaniel Goble, Ahmet S. Asan, Kira S. Brenner, Nate Shepard, Ana Geller, Katherine J. Stalnaker, Olivia Seidel, Yining Wang, Ally Nicolella, Bryan J. Song, Sean P. Moran, Hasmik Keshishian, Steven A. Carr, Jen Q. Pan, Morgan Sheng

**Author notes:** Correspondence (S.A.); (M.S.).

## Abstract

Rare loss-of-function variants in *SRRM2*, which encodes a nuclear speckle scaffold and splicing factor, are associated with schizophrenia and neurodevelopmental disorders. How *SRRM2* haploinsufficiency disrupts brain function is unknown. We find that *Srrm2^+/-^* mice exhibit (i) large-scale changes in gene expression in neuronal and glial cells, affecting DNA-binding-, synapse-, translation-, mitochondria-related pathways across multiple brain regions; (ii) alterations in splicing and/or abundance of multiple postsynaptic proteins, including reduction of the gamma isoform of SynGAP and elevation of its interactor, Agap3; and (iii) reduced oligodendrocyte proportions, particularly in striatum, accompanied by decreased expression of myelin-related mRNAs and proteins. Human iPSC-derived neurons deficient in *SRRM2* display conserved *AGAP3* splicing defects. Behaviorally, *Srrm2^+/-^* mice have reduced locomotor activity and impaired startle responses, and EEG recordings reveal reduced sleep spindles resembling humans with schizophrenia. Our findings identify specific synaptic changes, splicing dysregulation, and impaired myelination as mechanisms linking *SRRM2* haploinsufficiency to neuropsychiatric disease.

## INTRODUCTION

Schizophrenia (SCZ) is a common psychiatric disorder and a leading cause of disability in young adults ^1^. Large-scale exome- and targeted-sequencing studies of humans with SCZ have associated rare heterozygous protein-truncating variants in serine/arginine repetitive matrix 2 (*SRRM2*) with substantially elevated disease risk (odds ratio ∼9; combined *P*-value = 7.2×10^−7^)^2,3^. Exome sequencing of ∼31000 parent/offspring trios has also linked heterozygous loss-of-function (LoF) of *SRRM2* to a syndromic neurodevelopmental disorder that presents with mild intellectual disability (ID) and mild-to-moderate developmental delay (DD), most prominently in language acquisition ^4,5^.

A gene highly intolerant to LoF mutations (probability of LoF intolerance = 1) ^5^, *SRRM2* encodes a ∼300 kDa RNA-binding protein that regulates pre-mRNA splicing ^6–8^. In yeast, the ortholog of SRRM2 co-purifies with spliceosome components ^9^. In mammalian cells, SRRM2 primarily localizes to and acts as a core scaffold of nuclear speckles ^8,10,11^, which are irregularly shaped biomolecular condensates organize pre-mRNA splicing by acting as highly concentrated reservoirs of splicing machinery ^12^. Consequently, the physical proximity of genes to speckles influences the splicing efficiency of their pre-mRNAs ^13^.

SCZ’s genetic link to *SRRM2*—and hence to nuclear speckles and mRNA splicing—is particularly interesting because (i) large-scale transcriptomic analysis of human brain tissue has revealed an outsized impact on mRNA isoform and splicing regulation in individuals with SCZ, compared to those with bipolar disorder or autism ^14^, and (ii) in the human fetal cortex, genomic regions in proximity of nuclear speckles are enriched for SCZ common variant risk loci ^15^. These molecular findings, together with the human genetics association, support a mechanistic relationship between *SRRM2*, RNA splicing, and SCZ pathophysiology.

Little is known about the pathomechanisms of SCZ or the *SRRM2* LoF disorder. We reasoned that a comprehensive characterization of *Srrm2* heterozygous mutant mice—at transcriptomic, proteomic, neurophysiological, and behavioral levels—should provide insight into the neurobiological aberrations in this genetic animal model of SCZ and *SRRM2* LoF disorder. In this study, we performed bulk and single-nucleus RNA sequencing of multiple brain regions at different ages, synapse proteomics of the neocortex, and behavioral and electroencephalogram (EEG) analyses to uncover the molecular and neurobiological endophenotypes of *Srrm2* heterozygous LoF (*Srrm2^+/-^*) mice.

Combined, we find that haploinsufficiency of *Srrm2* in mice leads to impaired expression of synapse-related genes and key postsynaptic proteins, deficits in oligodendrocytes and myelination, and EEG abnormalities that resemble those seen in humans with SCZ. Our study highlights synapse dysfunction, abnormal splicing, and impaired myelination as potential mechanisms of SCZ and the *SRRM2* LoF neurodevelopmental disorder.

## RESULTS

### Large-scale and correlated transcriptomic changes across different brain regions in *Srrm2*^+/-^ mice

To evaluate the impact of *Srrm2* LoF on the brain in an unbiased manner, we used RNA sequencing (RNA-seq) to measure gene expression changes (compared to wild-type (WT) littermates) in 8 brain regions of *Srrm2*^+/-^ mice—the genotype linked to SCZ and the *SRRM2* LoF syndrome)—at 1- and 3-months of age (Fig. S1A). Principal component analysis showed clear clustering of bulk RNA-seq samples by brain region, consistent with accurate tissue dissection (Fig. S1A). The *Srrm2* gene itself was highly expressed across brain regions in WT animals (Fig. S1B), especially in the prefrontal cortex (PFC) and thalamus (TH). Expression of *Srrm2* was reduced in *Srrm2*^+/-^ mice—across the entire coding sequence (Fig. S1C)—to approximately 50% or less of WT in all brain regions (Fig. S1B).

Differential expression (DE) analysis of bulk-RNA-seq data revealed marked gene expression changes in all brain regions in *Srrm2*^+/-^ mice. At 1-month of age, *Srrm2*^+/-^ brains showed hundreds to thousands of DE genes (DEGs; defined as genes with FDR-adjusted *P* < 0.05 in DE analysis) in diverse brain regions (Fig. 1A), with the striatum (ST) showing the largest number of DEGs, followed by the cerebellar cortex (CC)(Fig. S1D). For every brain region there were many more DEGs at 1-month than at 3-months of age in *Srrm2*^+/-^ mice (Fig. 1A; Table S1).

**Fig. 1:**
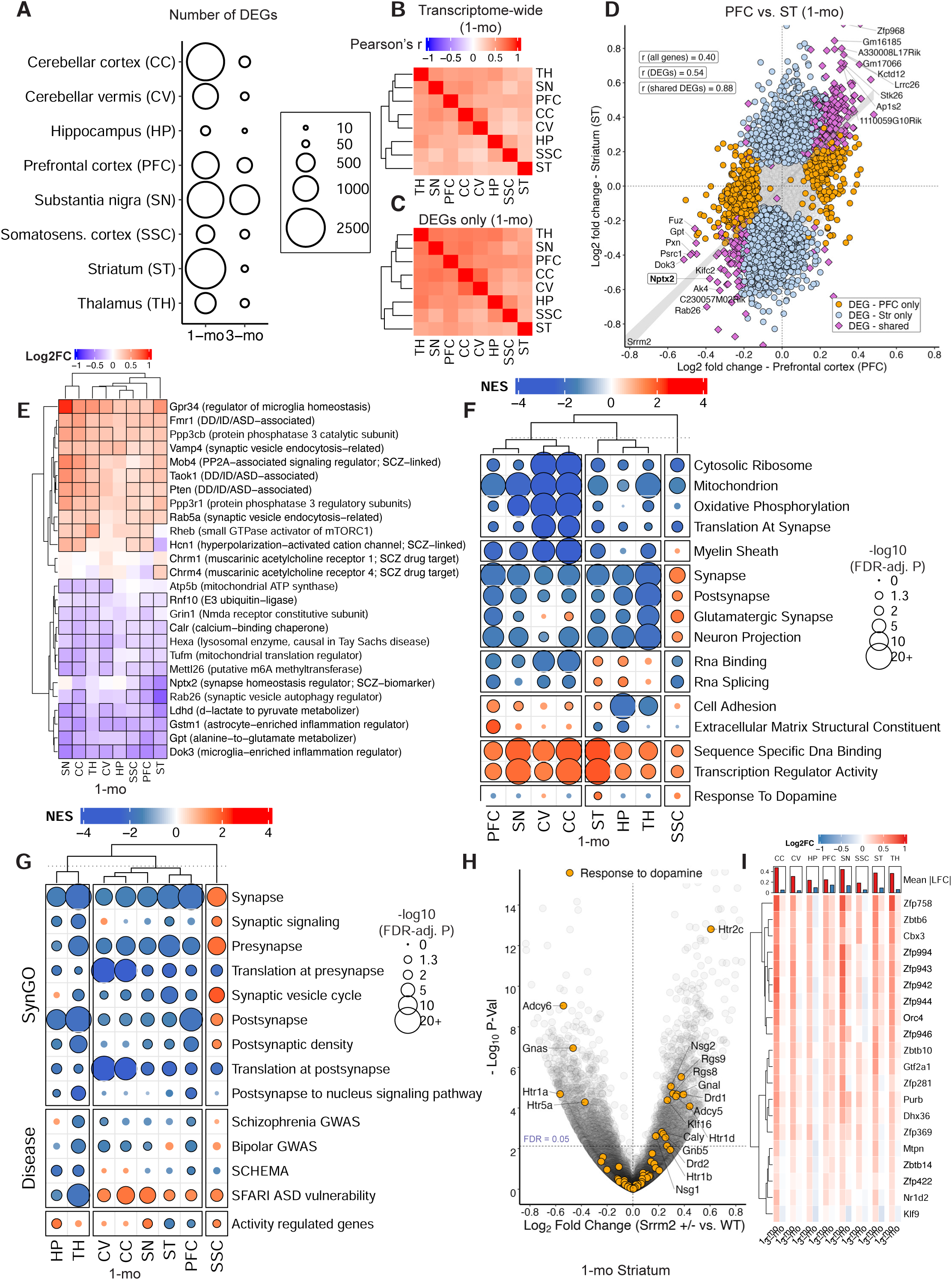
Large-scale and correlated transcriptomic changes across different brain regions in *Srrm2*+/- mice. **(A)** Number of DEGs (*Srrm2*+/- vs. WT) in the profiled brain regions at 1- and 3-months of age. RNA-seq libraries were prepared from 5 WT and 5 *Srrm2*+/- mice for each brain region and age, however three 3-month SN samples (1 WT and 2 *Srrm2*+/-) were excluded from sequencing due to failure in library preparation. Differential expression analysis was performed using DESeq2 (Wald test) with Benjamini–Hochberg FDR correction; DEGs are defined as genes with FDR-adjusted *P* < 0.05 in DE analysis. Correlation of gene expression changes (log2FC) in 1-month-old *Srrm2*+/- mice transcriptome-wide **(B)**, and **(C)** in the 6429 genes that are DE in any of the brain regions; correlations were calculated using Pearson’s correlation coefficient. **(D)** Evaluation of gene expression changes (log2FC) in the PFC against those in the striatum (ST) in 1-month-old *Srrm2*+/- mice. **(E)** Heatmap showing log2FCs of genes of interest that are DE in at least 4 brain regions in 1-month-old *Srrm2*+/- mice. Also shown are Chrm1/4 and Nptx2. Stroked cells indicate DE in the indicated brain region. GSEA was performed on genes ranked by the DESeq2 Wald test statistic; gene sets were considered significantly enriched at FDR-adjusted *P* < 0.05. GSEA results showing enrichment of various molecular pathways from **(F)** MSigDB M5 ontology gene sets, and **(G)** SynGO (v1.2) and curated neuropsychiatric disease and activity-related gene sets. **(H)** Evaluation of gene expression changes (log2FC) against *P*-value in the striatum of 1-month-old *Srrm2*+/- mice; genes in the “response to dopamine” pathway are highlighted. Horizontal dotted line indicates FDR-adjusted *P*-value threshold of 0.05. **(I)** Heatmap displaying log2 fold-changes for the top 20 leading-edge genes from the “Sequence-Specific DNA Binding” GO term. The top barplot indicates the mean absolute log2 fold-change for each column, highlighting the magnitude of regional transcriptional shifts.

By comparing log2 fold change (Log2FC) of all individual genes between different brain regions, we observed considerable similarity in transcriptomic changes across different brain regions, particularly in 1-month old *Srrm2*^+/-^ mice (Pearson’s r = 0.20–0.63, median r = 0.39, *P* < 2×10^-16^ in every comparison; Fig. 1B). Restricting this comparison to only the 6429 DEGs (FDR < 0.05 in at least one brain region) revealed even stronger correlations between different brain regions (Fig. 1C; r = 0.35–0.76, median r = 0.55).

Correspondingly, DEGs showed strong overlap between different brain regions (Fig. S1E). The 28 DEGs shared across all regions (Fig. S1F), and the 1299 DEGs shared between any two regions (Fig. S1G), showed consistent directional changes across brain regions. Scatterplot of the log2FCs of all genes in ST against those in PFC—two brain regions with very different cell type populations—also showed correlated changes (r = 0.40 for all genes, r = 0.54 for DEGs in either PFC or ST; r = 0.88 for DEGs in both PFC and ST; Fig. 1D). Of note, *Nptx2* (neuronal pentraxin 2, an activity-regulated synapse-related gene) was strongly downregulated in both ST and PFC bulk RNA-seq (Fig. 1D). The NPTX2 protein is reduced in the cerebrospinal fluid ^16^ and in PFC synaptic fractions ^17^ of humans with SCZ, and is also a CSF biomarker for cognitive impairment in Alzheimer’s disease ^18^.

Among DEGs shared across at least four brain regions of 1-month-old *Srrm2*^+/-^ mice (n=488; Table S1), we noted widespread reduction in *Grin1* (NMDA receptor obligatory subunit), *Rab26* (synaptic vesicle autophagy regulator), and *Gpt* (alanine aminotransferase, which metabolizes alanine to produce glutamate (Fig. 1E). *Fmr1*, *Pten*, *Taok1* (all DD/ID associated genes), and *Gpr34* (a marker of homeostatic microglia) were elevated across many regions (see Fig. 1E, which also shows other interesting DEGs with their functions annotated).

At 3 months, there were similar, albeit more modest, correlations in gene expression changes and overlap in DEGs across brain regions of *Srrm2*^+/-^ mice (Fig. S1H–J). The 26 genes differentially expressed in four or more brain regions in 3-month-old *Srrm2*^+/-^ mice also exhibited consistent changes across different brain regions (Fig. S1K). Gene expression changes at 1-month vs 3-month were also correlated in all brain regions (Pearson’s r = 0.14 (HP)–0.58 (PFC) transcriptome-wide, r = 0.15 (HP)–0.70 (PFC) for DEGs at either age; *P* < 2×10^-16^ in all comparisons) except in ST, which had limited correlation (r = -0.008 transcriptome-wide, r = 0.025 for DEGs at either age). In summary, *Srrm2*^+/-^ mice showed major gene expression changes that are substantially correlated across brain regions and between 1 and 3 months of age.

### Molecular pathways affected by *Srrm2* LoF

To gain insight into the molecular pathways that were perturbed with *Srrm2* LoF, we performed gene set enrichment analysis (GSEA) on the bulk RNA-seq data from all brain regions using mSigDb ^19^, and SynGO ^20^ gene ontology (GO) terms, focusing on 1-month age, when transcriptomic changes were greater (Fig. 1A). Consistent with the transcriptome-wide correlations in gene expression changes, we observed multiple GO terms in mSigDB that were commonly changed across most brain regions in 1-month-old *Srrm2*^+/-^ mice. Gene sets related to synapse/glutamatergic synapse, ribosomes/translation, mitochondria/respiration, and myelination (a gene set of particular interest, see below) were downregulated, while those related to DNA binding and transcription regulation were upregulated (Fig. 1F; Table S2).

Using SynGO annotated terms (Fig. 1G), we observed downregulation of both presynapse- and postsynapse-related gene sets in all brain regions of 1-month-old *Srrm2*^+/-^mice, except SSC, which showed modest upregulation of many synapse-related terms (Fig. 1G). GSEA using SynGO and mSigDb gene sets both highlighted downregulation of translation-related genes across all brain regions, especially the cerebellum (Fig. 1F, 1G).

Genes associated with bipolar disorder ^21^ and schizophrenia ^22^ by GWAS (or by SCHEMA ^2^) were enriched in the downregulated genes in most of the brain regions of *Srrm2*^+/-^mice, most significantly the thalamus (Fig. 1G). Meanwhile, ASD risk genes from the SFARI database were enriched in upregulated genes in most brain regions, but not the thalamus or hippocampus. Only in the thalamus was there concordant enrichment of ASD and SCZ and BP genes among downregulated genes (Fig. 1G).

Activity regulated genes—the expression of which can be used as a surrogate measure of neuronal activity—were enriched in downregulated genes in PFC and ST, and in upregulated genes in HP, SSC, SN (Fig. 1G), indicating differential effects on regional brain activity in *Srrm2* mutants.

Because elevated dopamine signaling in the striatum (ST) is implicated in the pathophysiology of SCZ ^23^, we were intrigued to find by GSEA that the “response to dopamine” gene set (mSigDB) was significantly upregulated at 1 month age specifically in the striatum of *Srrm2*^+/-^ mice (Fig. 1F). Closer inspection revealed that although most genes in this gene set tended to increase (in line with the GSEA result), some genes were reduced in expression (Fig. 1H). Interestingly, the genes that were increased in *Srrm2*^+/-^ striatum—including *Htr2c, Drd1*, and *Drd2* which encode the serotonin 2C- and dopamine D1- and D2-receptors, and other genes like *Adcy5*, *Rgs9*, *Rgs8*, *Nsg2*, *Gnal* (Fig. 1H)—are known to be highly enriched in spiny projection neurons (SPNs) of the striatum ^24^. In contrast, the genes in the “response to dopamine” set that were robustly reduced (*Adcy6*, *Gnas*, *Htr1a*, *Htr5a*) are selectively expressed in other cell types (such as inhibitory interneurons) of the striatum (Fig. 1H) ^24^. These data point to a perturbation of the striatal circuitry in *Srrm2*^+/-^ mice.

Transcriptomic differences between *Srrm2*^+/-^ and WT mice, as measured by DEG counts, are attenuated at 3 months relative to 1 month (Fig. 1A). To examine age-dependent effects, we performed an *Age × Genotype* interaction analysis across RNA-seq data from all eight brain regions (see Methods). This analysis revealed that genes and pathways related to DNA-binding and transcriptional regulation, which are elevated at 1 month across *Srrm2*^+/-^ brain regions (Fig. 1F), show markedly less upregulation at 3 month age (Fig. 1I, Fig. S1L). In contrast, ribosomal and mitochondrial pathways, which are downregulated at 1 month (Fig. 1F), exhibit increased expression by 3 months (Fig. S1L). Thus, distinct molecular pathways show attenuation or reversal of genotype effects with age, possibly reflecting compensatory processes in the *Srrm2*^+/-^ brain.

### *Srrm2* LoF causes isoform-specific changes in SynGAP and Agap3 at the synapse

GSEA indicated widespread transcriptomic changes in synapse-related pathways across all brain regions in *Srrm2*^+/-^ mice. To investigate synaptic changes at the protein level we performed mass spectrometry-based quantitative proteomics on synapse fractions purified from cerebral cortex of 1- and 4-month-old *Srrm2*^+/-^ mice.

We observed 35 and 44 differentially expressed proteins (DEPs; defined as *P*-value < 0.01) in 1- and 4-month-old *Srrm2*^+/-^ mice synapses, respectively (Fig. 2A, 2B; Table S2). GSEA of the synaptic proteome changes using SynGO gene sets indicated a prominent reduction of several protein sets related to postsynaptic structure and function in 4-month-old *Srrm2*^+/-^ mice (Fig. 2C, Table S2). *Srrm2*^+/-^ synapses showed reductions in glutamatergic signaling proteins, especially GTPase regulatory proteins, such as SynGAP (encoded by *Syngap1*) and its binding partners Iqsec2, Iqsec1 ^25^; the retrovirus-like activity-dependent intercellular signaling protein Arc; multiple glutamate receptor subunits and associated proteins (Gria3, Gria2, Olfm2, Grid2ip); and postsynaptic scaffolding and cytoskeletal proteins (Anks1b, Psd95 (encoded by *Dlg4*), Cript) (Fig. 2D, Fig. S2A). This downregulation of key postsynaptic proteins—reminiscent of synapse proteome changes in humans with SCZ ^17^—was less marked in 1-month-old *Srrm2*^+/-^synapses (Fig. S2B), indicating that these postsynaptic protein changes lag behind the transcriptomic changes in synapse pathways (Fig. 1F, 1G). Additionally, GSEA with mSigDB terms showed that protein sets related to RBPs, ribosomes, and translation were upregulated in the synapses of 4-month-old *Srrm2*^+/-^ mice; these same pathways were either downregulated or minimally changed at 1-month (Fig. 2C, Fig. S2C) further underscoring that the synapse proteome changes in an age-dependent manner in the *Srrm2*^+/-^ brain (Table S2).

**Fig. 2:**
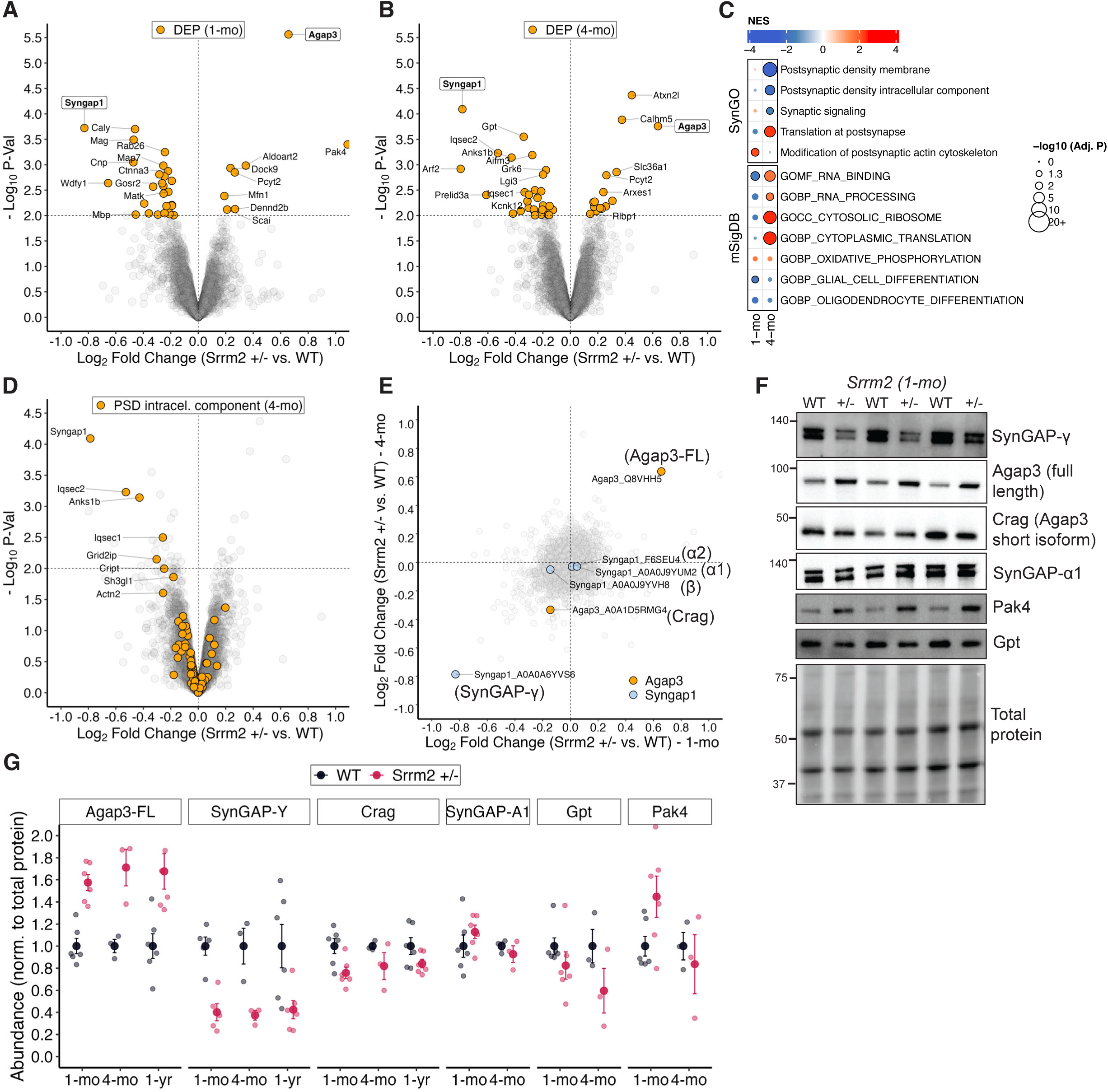
*Srrm2* LoF causes isoform-specific changes in SynGAP and Agap3 at the synapse. Volcano plots showing **(A)** DEPs in 1-month-old and **(B)** 4-month-old *Srrm2*+/- mice synapses (1-mo WT n = 4, 1-mo *Srrm2*+/-^-^ n = 5, 4-mo WT n = 5, 4-mo *Srrm2*+/- n = 4 animals). *Gene symbols* of the genes encoding some of the top DEPs are labeled. DEPs were assessed using moderated t-tests, and are defined as proteins with *P* < 0.01 in DE analysis. **(C)** GSEA results highlighting prominently changed pathways in SynGO and MSigDB M5 ontology sets in the cortical synapse proteome of *Srrm2*+/- mice. GSEA was performed using *fgsea* on preranked proteins (ranked by the moderated t-test statistic); gene sets were considered significantly enriched at FDR-adjusted *P* < 0.05. **(D)** Volcano plot of synapse proteome changes in 4-month-old *Srrm2*+/- mice, highlighting all proteins in the SynGO “postsynaptic density intracellular component” gene set. **(E)** Evaluation of log2FCs in 1-month-old *Srrm2*+/- mice synapses against those in 4-month-old *Srrm2*+/- mice; correlation was calculated using Pearson’s correlation coefficient. Protein isoforms of SynGAP and Agap3 are highlighted. **(F)** Western blots showing levels of SynGAP-γ, Agap3-FL, Crag, and SynGAP-A1, Pak4, and Gpt in synapse fractions from 1-month-old *Srrm2*+/- mice and WT littermates. **(G)** Quantifications of (F). n = 6 WT and 6 *Srrm2*+/- (1 month), 3 WT and 3 *Srrm2*+/- (3 months), 6 WT and 6 *Srrm2*+/- (1 year). Data are presented as mean ± SEM. No statistical tests were performed.

Among all DEPs, a SynGAP proteoform showed the largest drop in protein level (log2FC = ∼ -0.8) and the most significant reduction (by *P*-value) in *Srrm2*^+/-^ synapse fractions at both 1- and 4-months of age (Fig. 2A, 2B). This finding is especially interesting because *Syngap1* haploinsufficiency is a common cause of DD, as well as of ID and autism, all of which are observed in humans with *SRRM2* LoF mutations ^5,26^. Given the large reduction of SynGAP, it was also striking to find that Agap3—a known binding partner of SynGAP that is thought to regulate synaptic Ras/Erk signaling and endocytosis of AMPA receptors ^27–29^—was among the two most upregulated proteins (log2FC = ∼ 0.7) in *Srrm2*^+/-^ synapses at both ages (Fig. 2A, 2B).

SynGAP exists in four isoforms (α1, α2, β, and γ) due to alternative splicing of the 3’ terminus of *Syngap1* (Fig. S2E) ^30,31^, while two isoforms of Agap3 have been described—full length (Agap3-FL) and an N-terminally truncated isoform, called Crag (Fig. S2F) ^27^. Detailed analysis of MS/MS spectra of peptides revealed that in *Srrm2*^+/-^ synapses at both ages, it was specifically the gamma isoform of SynGAP (SynGAP-γ) that was diminished, and specifically the Agap3-FL isoform that was increased (Fig. 2E).

By western blotting of cortical synaptic fractions from an independent cohort of *Srrm2*^+/-^mice, we confirmed the changes in Agap3 and SynGAP isoforms, as well as in Gpt (alanine aminotransferase, which metabolizes alanine to produce glutamate), whose levels declined in line with its reduction in RNA expression (Fig. 1E), and in Pak4, a kinase that changed in an age-specific manner (Fig. 2F, 2G, Fig. S2D). The ∼60-70% increase in Agap3-FL and ∼60-70% reduction in SynGAP-γ was also seen at 1 year of age (Fig. 2G, Fig. S2D), indicating that this particular biochemical phenotype persists into late adulthood in *Srrm2*^+/-^ mice.

### *SRRM2* deficiency causes abnormal splicing of *AGAP3* mRNA across species

There were no major changes in *Agap3* and *Syngap1* total mRNA expression as measured by standard gene-level RNA-seq analysis (Fig. S3A). However, isoform-specific alterations may be masked when gene transcripts are considered in aggregate. We therefore investigated mRNA isoform abundance changes genome-wide by meta-analyzing *P*-values of changes of all the isoforms of a given gene (derived from differential isoform expression analysis) to obtain a gene-level *P*-value for every gene (see Methods and Yi et al. ^32^). Using this approach, we observed *Agap3* to be among the most significant DEGs in all brain regions in both 1- and 3-month-old *Srrm2*^+/-^ mice (Fig. 3A; Table S3). The majority of the six annotated mRNA isoforms of *Agap3* (*201* through *206*) were changed with low *P*-values in every brain region at both ages (Fig. S3B), and the aggregation of the *P*-values of the individual isoforms resulted in highly significant *P*-values for the *Agap3* gene (Fig. 3A).

**Fig. 3:**
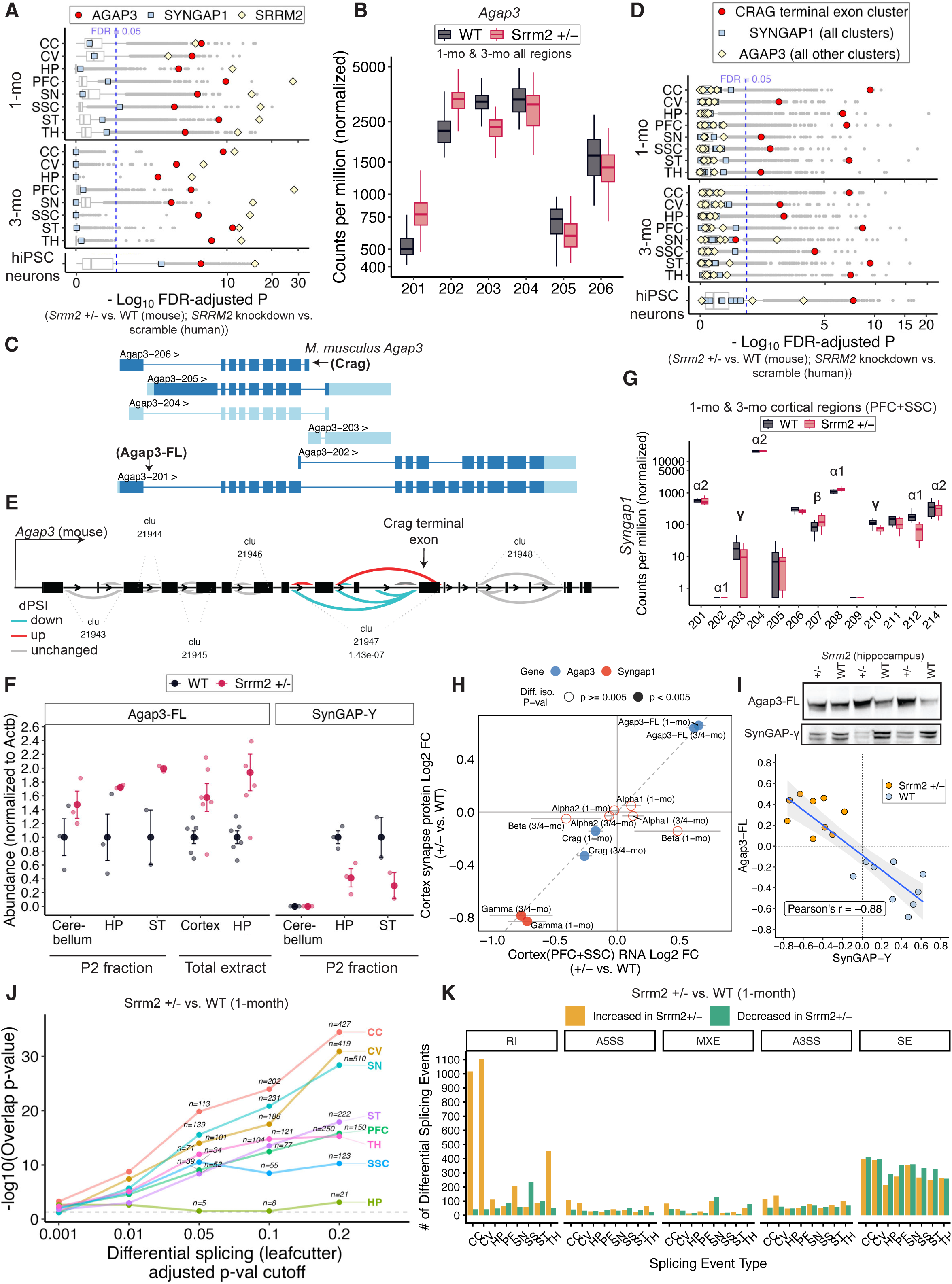
*SRRM2* deficiency causes abnormal splicing of *Agap3*. **(A)** Each boxplot shows distributions of gene-level *P*-values of transcriptomic differences (*Srrm2*+/- vs. WT) in the indicated brain region and age, as well as those observed with shRNA knockdown of *SRRM2* (vs. scramble shRNA) in human iPSC-derived glutamatergic neurons. Isoform-level differential expression was performed using *Sleuth*; gene-level *P*-values were obtained by aggregating isoform-level *P*-values using the Lancaster method. Human iPSC RNA-seq data were obtained from Wu (2024) ^33^. Counts of individual mRNA isoforms of *Agap3* across all 8 brain regions in 1- and 3-month-old *Srrm2*+/- mice and WT littermates **(B)**, and gene models showing exonic and intronic regions of the six Ensembl-annotated *Mus musculus Agap3* isoforms **(C)**. Distributions of intron-cluster *P*-values from Leafcutter splicing analysis in 1- and 3-month-old *Srrm2*+/- mice and in *SRRM2* shRNA-treated human iPSC-derived glutamatergic neurons **(D)**. Differential splicing was assessed using Leafcutter (likelihood ratio test with FDR correction). **(E)** Gene model highlighting splicing changes in *Agap3* identified by Leafcutter. “Clu” and associated dotted lines indicate intron clusters, and colors indicate changes in percent spliced in (PSI). **(F)** Quantification of Agap3 and SynGAP-γ protein levels in the P2 fractions and total extracts of the indicated brain regions of *Srrm2*+/- mice and WT littermates. For P2 fractions, n = 3 WT and 3 *Srrm2*+/- for all regions except striatum (ST), where n = 2 WT and 2 *Srrm2*+/-. For total extracts, n = 6 WT and 6 *Srrm2*+/-. No statistical tests were performed. **(G)** Counts of individual mRNA isoforms of *Syngap1* in the cortex (PFC + SSC) in 1- and 3-month-old *Srrm2*+/- mice and WT littermates. The corresponding SynGAP protein isoforms (α1, α2, β, and γ) are labeled for reference; multiple *Syngap1* mRNA isoforms can encode the same protein isoform due to alternative transcription start sites and shared C-terminal splice architecture. **(H)** Evaluation of log2FC of SynGAP- and Agap3-encoding RNA isoforms relative to corresponding protein log2FC measured by cortical synapse proteomics (MS/MS). Data include age-matched 1-month RNA and proteomics datasets, as well as 3-month RNA compared to 4-month proteomics (labeled “3/4mo”). Horizontal error bars indicate DESeq2-derived log2FC standard errors. *P*-values represent nominal DESeq2 Wald test *P*-values for isoform-level differential expression. **(I)** Top: Representative western blot of Agap3-FL and SynGAP-γ in P2 fractions of the *Srrm2*+/-hippocampus. Bottom: Evaluation of MS/MS abundances of Agap3-FL compared to those of SynGAP-γ. **(J)** Overrepresentation analysis comparing DEGs (FDR-adjusted *P* < 0.05, Lancaster aggregation approach) with Leafcutter differentially spliced genes across a range of differential splicing significance thresholds. The y-axis denotes –log10 *P*-value of the statistical significance of overlap, which was determined using two-sided Fisher’s exact test. For the 0.05, 0.1, and 0.2 thresholds, labels indicate the absolute number of overlapping genes. **(K)** Number of different types of alternative splicing events regulated by *Srrm2* LoF across brain regions, as found by rMATS. SE, skipped exon; A5SS, alternative 5′ splice site; A3SS, alternative 3′ splice site; MXE, mutually exclusive exon; RI, retained intron. Error bars indicate SEM throughout.

Among the individual mRNA isoforms of *Agap3*, those that encompassed the 3’ end of *Agap3* were upregulated in the brains of *Srrm2*^+/-^ mice (by ∼60% for *Agap3-201,* which encodes the canonical Agap3-FL; Fig. 3B, 3C). Conversely, isoforms that lacked exons towards the 3’ end (including *Agap3-206*, annotated as encoding Crag) were downregulated (Fig. 3B, 3C). These findings imply that *Agap3* splicing may be altered with *Srrm2* LoF. Indeed, annotation-free examination of local splicing changes revealed that a singular intron cluster of *Agap3* (among 6 clusters) was prominently differentially spliced in every brain region of 1-month-old and 7 of 8 regions of 3-month-old *Srrm2*^+/-^ mice (Fig. 3D; Table S3); this complex splicing alteration leads to increased skipping of the terminal exon of Crag that also harbors its stop codon (Fig. 3E).

By re-analyzing publicly available data ^33^, we found that knockdown of *SRRM2* in human iPSC-derived glutamatergic neurons causes remarkably similar isoform and splicing changes in *AGAP3*. Specifically, we observe highly significant gene-level *P*-values for *AGAP3* (as detected by aggregation of *P*-values of isoform changes; Fig. 3A), elevation of isoforms encompassing its 3’ end (including *AGAP3-202,* which encodes full-length AGAP3; Fig. S3E, S3F), downregulation of transcripts encoding CRAG (Fig. S3E, S3F), and a robust splicing change (Fig. 3D) that leads to increased skipping of the terminal exon of CRAG (Fig. S3G, S3H). *SRRM2* loss therefore causes conserved splicing changes in *AGAP3*.

These findings demonstrate that *Srrm2* LoF favors splicing of *Agap3* towards increased production of its full-length mRNA isoform (which encodes Agap3-FL). Consistent with this mechanism, Agap3-FL protein was markedly elevated (∼60-95%) in P2 fractions (see Methods) of the cerebellum, hippocampus, and striatum, as well as in total extracts of the cortex and hippocampus (Fig. 3F, Fig. S3D).

In contrast to *Agap3*, *Syngap1* was notably unchanged in *Srrm2*^+/-^ mice in terms of mRNA splicing (Fig. 3D) and isoform-aggregation informed gene significance (Fig. 3A). Nonetheless, western blot examination of the same P2 fractions for SynGAP-γ protein—SynGAP-γ is not readily detectable in total brain extracts—showed ∼60-70% reduction of SynGAP-γ in *Srrm2*^+/-^ hippocampus and striatum (Fig. 3F, Fig. S3D; SynGAP-γ was not detected in cerebellum).

This finding prompted a higher-resolution transcriptomic re-evaluation focused on *Syngap1* (see Methods). Because γ-encoding *Syngap1* transcripts are expressed at low levels (∼10–100 counts per million), we pooled RNA-seq data from cortical regions (PFC + SSC) to increase statistical power for isoform analyses. Examination of *Syngap1* mRNA isoform counts revealed that the SynGAP-γ-encoding *Syngap1* transcripts (*203* and *210*) were reduced in *Srrm2*^+/-^ cortex (Fig. 3G). Differential isoform analysis of the pooled cortical RNA-seq data further revealed a combined log2FC of ∼ −0.8 for *Syngap1 203* and *210*—which arise from alternative start sites but encode the same SynGAP-γ C-terminus—closely matching the ∼ −0.8 log2FC depletion of SynGAP-γ protein observed in cortical synapse fractions (Fig. 3H). Together, these results indicate that *Srrm2* LoF selectively downregulates SynGAP-γ through reduced expression of the mRNA isoforms that encode SynGAP-γ.

Additionally, western blots of Agap3 and SynGAP-γ in individual animals showed that a large increase in Agap3-FL was associated with a large reduction in SynGAP-γ (Fig. 3I). Comparison of the MS/MS abundances of Agap3-FL against that of SynGAP-γ in the synapse proteomics confirmed a striking negative correlation between these two proteins across many individual mice (Fig. 3I). Combined with the known interaction between Agap3 and SynGAP, our findings establish that Srrm2 functions as a key orchestrator of a coordinated RNA regulation program that maintains the stoichiometry of the Agap3/SynGAP complex at the synapse.

Large-scale changes were also observed in the expression of mRNA isoforms across many genes (Fig. 3A), consistent with widespread splicing disruption in *Srrm2* mutant brains (Fig. 3D). To assess how these alterations relate to differential gene expression, we compared DEGs identified by gene-level aggregation of isoform change *P*-values (FDR < 0.05; Fig. 3A) with differentially spliced genes (Fig. 3D) defined across progressively relaxed significance thresholds. For nearly all brain regions, the significance of overlap increased monotonically as thresholds were relaxed, indicating that not only the strongest but also numerous lower-magnitude splicing alterations are non-randomly associated with differential expression (Fig. 3J). This progressive increase supports a model in which transcriptome alteration in *Srrm2*^+/-^brains arises from a distributed, system-wide loss of splicing fidelity, alongside high-effect alterations such as those observed for *Agap3*.

What is the overall nature of this splicing infidelity? rMATS analysis revealed widespread dysregulation across brain regions, dominated by intron retention (RI) and skipped exon (SE) events. *Srrm2* LoF triggered a surge in RI, most notably in the cerebellar regions, where ∼1,100 events were elevated compared to only ∼40 reduced (Fig. 3K; Table S3). SE events were also frequent across regions and showed a more balanced distribution; in most regions, events showing reduced skipping (increased inclusion) nevertheless remained slightly more prevalent than those showing elevated skipping. Notably, human iPSC-derived neurons with *SRRM2* knockdown ^33^ (alongside other human and mouse cell models ^11,34^) exhibit a similar predominance of RI and SE abnormalities, indicating that the pattern of splicing disruption caused by loss of *SRRM2* is conserved between humans and mice.

In sum, *Srrm2* LoF broadly compromises alternative splicing, thereby extensively remodeling the brain transcriptome, including pronounced, conserved effects on *Agap3* and *Syngap1*.

### Decreased oligodendrocyte and increased SPN proportions in the *Srrm2*^+/-^ brain

To examine the impact of *Srrm2* LoF on specific cell types of the brain, we carried out single-nucleus RNA-seq, focusing on the striatum and the prefrontal cortex (PFC) of 1-month-old *Srrm2*^+/-^ mice. Cell types were inferred using UMAP-based unsupervised clustering and annotated based on expression of marker genes (Fig. 4A).

**Fig. 4:**
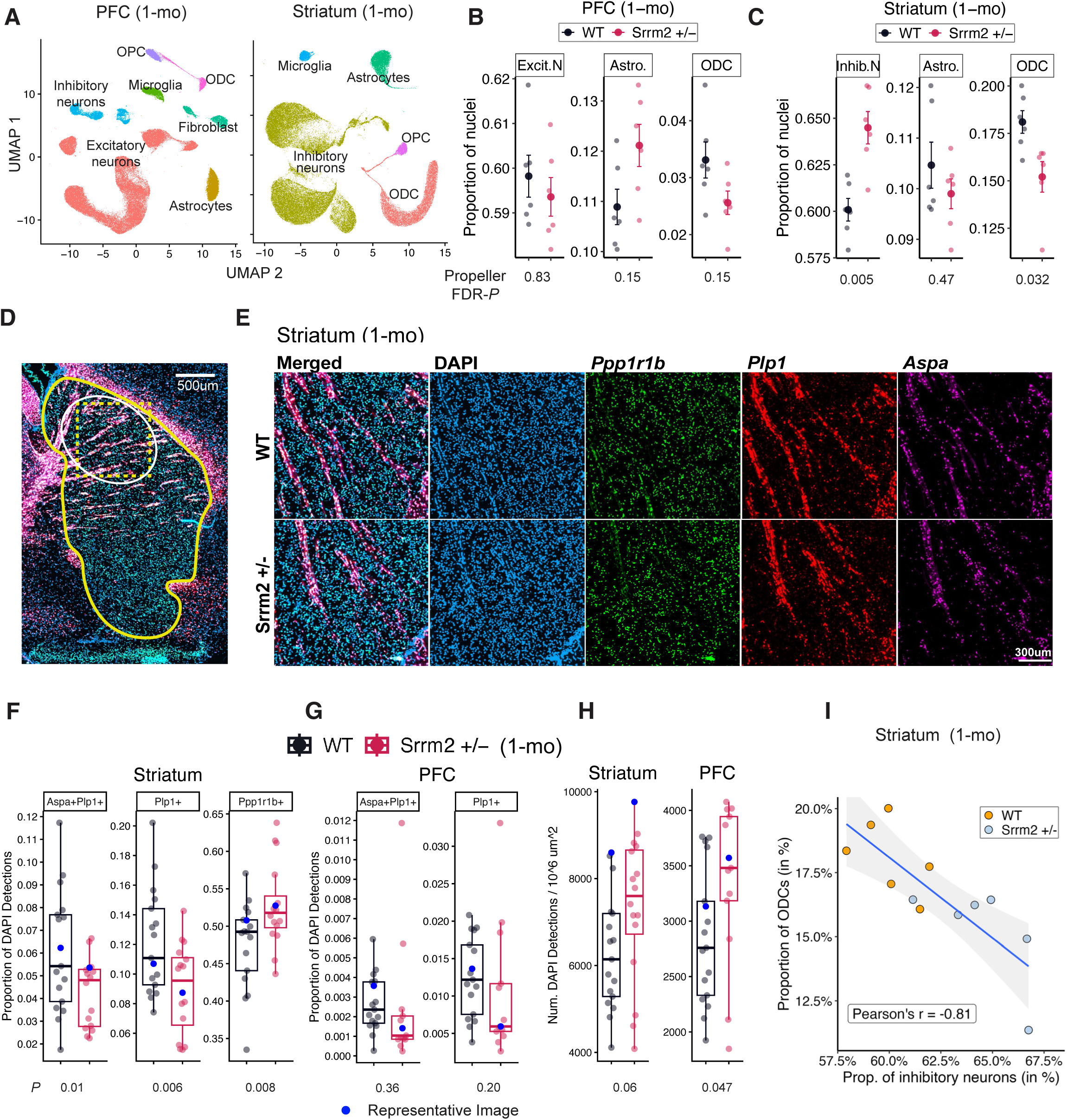
Decreased oligodendrocyte and increased SPN proportions in the *Srrm2*+/- brain. **(A)** UMAP representation of the major cell types identified by snRNA-seq in the indicated brain regions of WT and *Srrm2*+/- mice. snRNA-seq libraries were prepared from 6 WT and 6 *Srrm2*+/- mice for each brain region. Cell type proportions of nuclei profiled from **(B)** PFC and **(C)** striatum of 1-month-old *Srrm2*+/-mice and WT littermate controls (Inhib.N = inhibitory neurons, Excit.N = excitatory neurons, Astro. = astrocytes, ODC = oligodendrocytes). Statistical testing for differences in cell type proportions between genotypes was performed using the propeller function in speckle with arcsin normalization and Benjamini–Hochberg FDR correction. Error bars indicate SEM. **(D)** Representative sagittal brain section highlighting the striatum area (yellow outline) (10 μm) of *Srrm2*+/- mice probed by RNAScope. The white oval denotes the data collection region; the dotted square indicates the high magnification areas shown in (E). Because the dorsal striatum was the region dissected for both bulk RNA-seq and single-nucleus RNA-seq, RNAscope quantification was restricted to the dorsal striatum. **(E)** Representative images showing DAPI (nuclei), *Ppp1r1b* (SPNs), *Plp1* (differentiated oligodendrocytes), and *Aspa* (mature oligodendrocytes) expression in WT vs *Srrm2*+/- striatum. Data points corresponding to these specific images are highlighted in blue in **(F–H)**. See Fig. S4 for additional images. Boxplots for **(F)** striatum and **(G)** PFC showing the proportion of *Plp1*□/*Aspa*□ mature oligodendrocytes, *Plp1*□ pre-myelinating and mature oligodendrocyte cells, and *Ppp1r1b*□ D1 and D2 SPNs relative to total number of DAPI detections. *P*-values were determined using a one-tailed Student’s t-test. **(H)** Boxplots showing the number of DAPI+ cells normalized by area. *P*-values were calculated using a two-tailed Student’s t-test. For **(F–H)**, data represent 6 WT and 6 *Srrm2*+/- littermates (all males) at P29 with three sections per animal; each point = 1 section. **(I)** snRNA-seq evaluation of inhibitory neuron vs. ODC proportions (in percentages) in 1-month striatum of *Srrm2*+/- mice and WT littermates. Correlation was assessed using Pearson’s r.

Cell composition analysis of snRNA-seq data revealed a reduction in proportion of oligodendrocytes (ODCs) across PFC (reduced by ∼23%; FDR-adjusted *P* = 0.15) and striatum (by ∼16%; FDR-adjusted *P* = 0.03) of *Srrm2*^+/-^ mice (Fig. 4B, 4C; Table S4). We validated this finding *in situ;* by RNAscope we observed reduced proportions (number of cells / total number of DAPI labeled cells) of *Plp1*lJ (premyelinating and mature oligodendrocytes) and *Plp1*lJ*/Aspa*lJ cells (mature oligodendrocytes) in the striatum of *Srrm2*^+/-^ mice (*Plp1*lJ: - ∼25%, *P* = 0.006; *Plp1*lJ*/Aspa*lJ: - ∼29%, *P =* 0.01) (Fig. 4D-F; Table S4). In the PFC, ODC proportions were also reduced, although these differences did not reach statistical significance (*Plp1*lJ: - ∼18%; *Plp1*lJ*/Aspa*lJ: - ∼12%) (Fig. 4G, Fig. S4B-E; Table S4). The greater variability in the PFC likely reflects the much lower abundance of ODCs in this brain region (∼10-fold fewer than in striatum), which limits power to detect modest changes.

In contrast to the reduction in oligodendrocytes, the proportion of inhibitory neurons in the striatum rose by ∼7% (FDR-adjusted *P* = 0.005; Fig. 4C). This change was driven by a ∼7–8% elevation in both direct (D1)- and indirect (D2)- pathway SPNs (Fig. S4A). RNAscope analysis confirmed ∼10% increase (*P =* 0.01) in *Ppp1r1b+* cells (a marker of SPNs) in the striatum of *Srrm2*^+/-^ mice (Fig. 4F). The rise in SPN proportions is interesting because this increase may explain, in part, the increased expression of SPN-specific genes in striatum bulk RNA-seq (see above, and Fig. 1H). Notably, lineage expansion of SPNs has also been reported in the 16p11.2 microdeletion mouse model of neurodevelopmental disorders ^35,36^.

RNAscope further revealed an increase in total cell density (measured by DAPI+ cells /area) in both the PFC (∼18%. *P =* 0.05) and striatum (∼16%, *P =* 0.06) of *Srrm2*^+/-^ mice (Fig. 4H). This finding is consistent with the observed cell composition shifts in *Srrm2*^+/-^ brains, in which specific populations—such as SPNs—increase in absolute abundance (cells/area), whereas oligodendrocytes are reduced as a fraction of total DAPI+ cells.

In humans with SCZ, there is substantial evidence for both a widespread decline in ODCs (including in the striatum) and the dysfunction of dopamine signaling ^37,38^. We observed a striking negative correlation between the proportion of ODCs and SPNs (the major recipients of dopamine signaling) in the striatum across individual WT and *Srrm2*^+/-^ mice, using both sn-Seq (Fig. 4I) and *in-situ* approaches (Fig. S4F). Thus, dopamine-responsive neurons dominate at the proportional expense of myelinating glia in the *Srrm2*^+/-^ striatum.

### Decrease in myelin-related mRNAs and proteins in *Srrm2*^+/-^ mice

Consistent with the reduction in proportion of ODCs (Fig. 4B, 4C), the PFC and striatum of 1-month *Srrm2*^+/-^ mice also showed significant downregulation of the myelin sheath gene set by GSEA of bulk RNA-seq data (Fig. 1F). In fact, the myelin sheath gene set was downregulated in most of the brain regions at both 1- and 3-months of age in *Srrm2*^+/-^ mice (Fig. 5A, Fig. 1F). Similarly, an ODC marker gene set (containing for ex. *Mal*, *Mog*, *Myrf*, *Cnp*, *Olig2*) was enriched in downregulated genes in most brain regions at both ages (Fig. S5A).

**Fig. 5:**
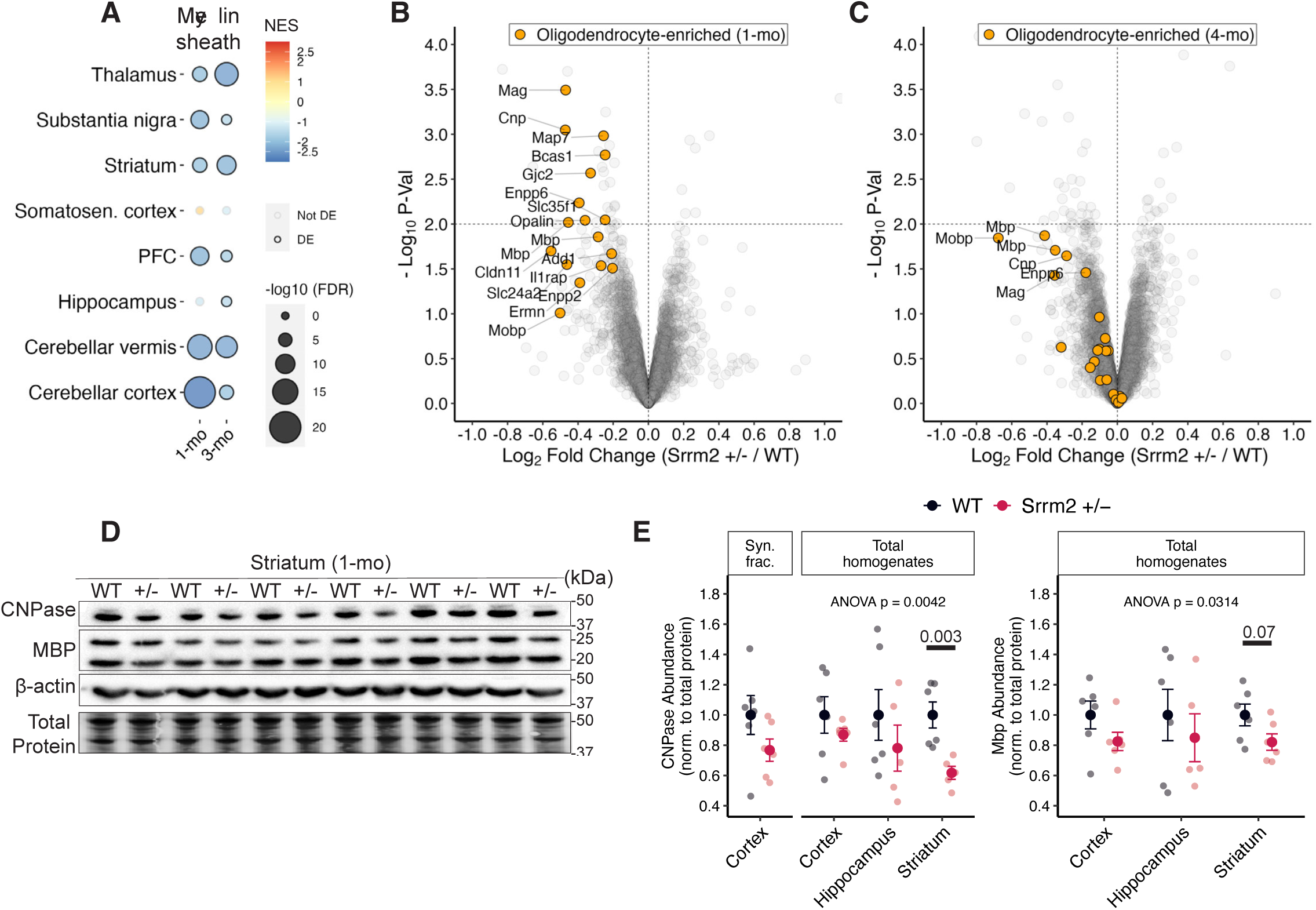
Decrease in myelin-related mRNAs and proteins in *Srrm2*+/- mice. **(A)** Bubble plot showing bulk RNA-seq GSEA results for the “myelin sheath” gene set in 8 brain regions of 1- and 3-mo *Srrm2*+/- animals. GSEA was performed on preranked genes (ranked by the DESeq2 Wald test statistic); gene sets were considered significantly enriched at FDR-adjusted *P* < 0.05. **(B)** Evaluation of cortical synapse proteome changes in 1-month-old *Srrm2*+/- mice (WT n = 4, *Srrm2*+/- n = 5 animals). DE was assessed using moderated t-tests. Downregulated proteins with P < 0.05 that were also found to show ODC enrichment in Allen Brain Atlas are highlighted. Also highlighted is Mobp. **(C)** Volcano plot of cortical synapse proteome changes in 4-month-old *Srrm2*+/- mice. The same set of proteins highlighted in (B) are highlighted. **(D)** Western blot of myelin markers Mbp and CNPase in striatal total extracts of WT and *Srrm2*+/- mice (n = 6 each). A representative section of total protein (the loading control) and an unchanged protein (β-Actin) is also shown. **(E)** Quantification of western blots examining changes in CNPase and Mbp in total homogenates or synapse fractions (Syn. frac.) in the indicated brain regions. Data are presented as mean ± SEM. Two-way ANOVA F-test *P*-value testing for difference in overall Mbp or CNPase levels between WT and *Srrm2*+/- across all examined brain regions is shown. Also shown are two-tailed t-test *P*-values testing for differences in either protein in the striatum.

While substantially depleted during purification of synapses, myelin is nonetheless present in synapse fractions ^39^; myelin proteins therefore can be reliably quantified in synapse fractions with sensitive approaches like mass-spectrometry. We noted in our synapse proteomics data that gene/protein sets related to oligodendrocyte differentiation were enriched among downregulated proteins in *Srrm2*^+/-^ synapse fractions (Fig. 2C). Remarkably, about a third (10/27) of downregulated DEPs in 1-month-old *Srrm2*^+/-^ synapse fractions (Fig. S5B) were either exclusively expressed or highly enriched in ODCs (Fig. 5B) ^40^. Most of the DEPs that are expressed primarily in ODCs (ex. CNPase (encoded by *Cnp*), Mbp, Mag, Bcas1, Opalin, Gjc2) are known to be specific components of the myelin sheath. The same set of ODC/myelin-associated proteins were found also at lower levels in 4-month-old *Srrm2*^+/-^ synapses, including key myelin proteins Plp, CNPase, Mbp, Mag, although the changes did not reach significance for any individual protein (Fig. 5C).

We validated the proteomics findings by western blotting of *total* lysates of cortex, striatum, and hippocampus, which showed ∼15-40% reduction of Mbp and CNPase protein levels in these brain regions of 1-month-old *Srrm2*^+/-^ mice, with particularly robust reduction in the striatum (Fig. 5D, 5E). We also confirmed by western blotting that CNPase was reduced ∼25% in synapse fractions purified from the cortex of an independent cohort of 1-month-old *Srrm2*^+/-^ mice (Fig. 5E). Thus, myelin proteins are downregulated in total extracts of *Srrm2*^+/-^ mouse brain; this likely accounts for the similar degree of reduction of myelin proteins in synaptic fractions. Together with the reduced ODC densities, these biochemical findings suggest myelination may be impaired in the brains of *Srrm2*^+/-^ mice. In this context, we note that *Qki*, which encodes an RBP that functions as a key regulator of ODC development ^41^, is among the most prominent DEGs—as measured by aggregation of *P*-values of isoform changes—across all brain regions of 1-month-old *Srrm2*^+/-^ mice (Table S3).

### *Srrm2* LoF causes pervasive gene expression changes in neuronal and glial cells

By DE analysis of the snRNA-seq data, 1-month-old *Srrm2*^+/-^ mice exhibited >1000 unique DEGs (defined as genes with FDR-adjusted *P* < 0.01 in DE analysis) aggregated across all cell types in the PFC, and ∼500 unique DEGs aggregated across all cell types in the striatum (Fig. 6A; Table S5). Transcriptomic changes were observed in all subtypes of neurons in both regions (Fig. 6A), with the largest number of DEGs seen in excitatory neurons of the PFC (especially layer 2/3 and 6 (L2/3IT and L6IT), and in inhibitory neurons of the striatum (direct-and indirect- SPNs) (Fig. 6A). Non-neuronal cells also showed sizable snRNA-seq changes, most prominently in astrocytes in the PFC and ODCs in the striatum (correlating with the impact on astrocyte and ODC proportions in the PFC and striatum, respectively (Fig. 4B, 4C)). Thus, losing one copy of *Srrm2* causes extensive changes in snRNA-seq in neuronal and glial cells in mice.

**Fig. 6:**
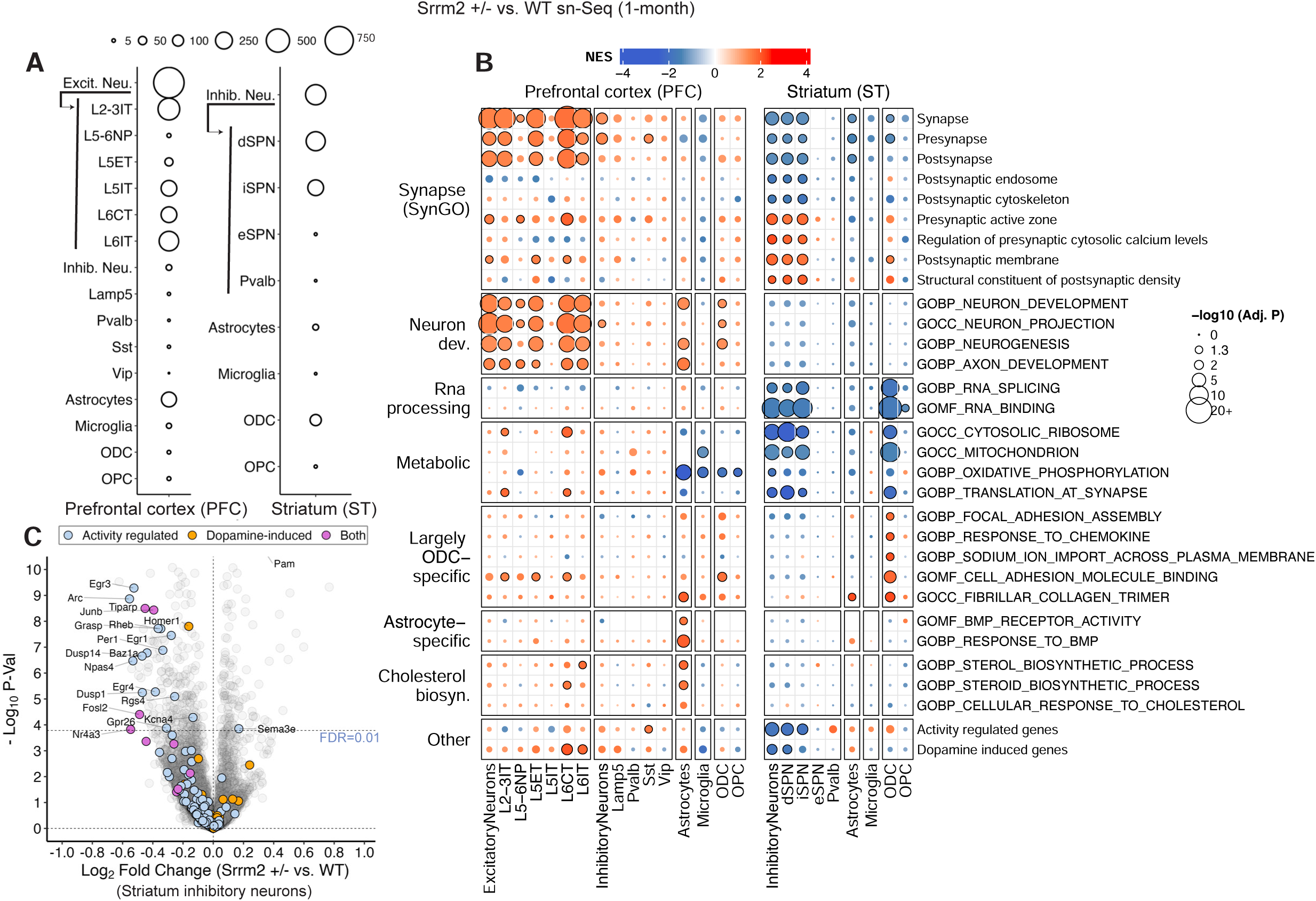
*Srrm2* LoF causes pervasive gene expression changes in neuronal and glial cells. **(A)** Number of DEGs across cell types in the PFC and striatum of 1-month-old *Srrm2*+/- mice. snRNA-seq libraries were prepared from 6 WT and 6 *Srrm2*+/- mice for each brain region. DEGs are defined as genes with FDR-adjusted *P* < 0.01 in differential expression analysis (Methods). **(B)** Molecular pathway changes as measured by GSEA across cell types in PFC and striatum of 1-month-old *Srrm2*+/- mice. GSEA was performed on preranked genes (ranked by the moderated t-test statistic); gene sets were considered significantly enriched at FDR-adjusted *P* < 0.05. **(C)** Evaluation of log2FC vs. *P*-value in inhibitory neurons of the striatum of 1-month-old *Srrm2*+/- mice; the genes that constitute activity-regulated genes and dopamine-induced genes, as well as those that belong to both, are highlighted. Horizontal dotted line indicates FDR = 0.01.

GSEA of the snRNA-seq changes revealed that synapse- and neuron development-related terms were upregulated in most excitatory neuron-types of the PFC in *Srrm2*^+/-^ mice (Fig. 6B; Table S5). Inhibitory neurons (dSPNs, iSPNs) of the striatum in *Srrm2*^+/-^ mice exhibited upregulation of some specific gene sets related to pre- and postsynaptic structure and function (e.g. presynaptic active zone, postsynaptic membrane), but the wider synapse sets (such as Synapse, Presynapse, Postynapse) were downregulated (Fig. 6B). The latter result in SPNs may be driven by downregulation of postsynaptic endosome and postsynaptic cytoskeleton gene sets, which did not occur in PFC excitatory neurons (Fig. 6B).

In glial cells, several gene sets, including those related to cell-adhesion (*Ctnna3*, *Cdh19*, *Itgad*), focal adhesion assembly (*Ptpra, Fmn1*), sodium ion import (*Slc5a6*, *Slc8a1*, *Slc6a1*), fibrillar collagen (*Col11a2*, *Col27a1*), and chemokine response (*Robo1*, *Stk39*) were upregulated in ODCs, while terms related to bone morphogenetic protein (BMP) receptor activity (*Chrdl1*, *Bmpr1b*) were downregulated in PFC astrocytes (Fig. 6B). Pathways related to sterol/cholesterol biosynthesis— reported to be downregulated in postmortem PFC astrocytes of humans with SCZ ^42^—were significantly upregulated in astrocytes of PFC but not significantly changed in striatum in *Srrm2*^+/-^ mice.

Expression of many activity-regulated genes, useful as surrogate of neuronal activity, was significantly reduced in SPNs of *Srrm2*^+/-^ mice (Fig. 6C), consistent with GSEA analysis of the activity-regulated gene set in both direct and indirect SPNs (Fig. 6B). Dopamine-induced genes, which overlap somewhat with the activity-regulated gene set, were also downregulated in SPNs (Fig. 6B, 6C). Overall, these snRNA-seq changes suggest that neuronal activity and/or dopamine signaling may be diminished on a per cell basis in *Srrm2*^+/-^ SPNs, the main neuron-type of the striatum. However, we note that the number and proportion of D1 and D2 SPNs went up in striatum (see above, Fig. 4). Interestingly, dopamine-induced genes were upregulated in excitatory neurons of PFC, particularly in L6CT and LCIT neurons (Fig. 6B).

Correlated with their reduced expression of activity-regulated genes, striatal SPNs of *Srrm2*^+/-^ mice showed downregulation of certain metabolic pathways (ribosome, protein synthesis, mitochondria, oxidative phosphorylation; Fig. 6B). RNA binding and RNA splicing-related gene sets (including *Son*, whose protein product forms a core scaffold of nuclear speckles with Srrm2), were also downregulated in SPNs (as well as in ODCs) of striatum, but were unchanged in PFC neurons (Fig. 6B).

### Behavioral abnormalities in *Srrm2*^+/-^ mice

We evaluated young adult *Srrm2*^+/-^ mice (4-5 months age) and their WT littermates in a battery of behavioral assays. *Srrm2*^+/-^ mutant mice of either sex showed no significant difference from WT in novel object recognition (Fig. S6A, S6B), 3-chamber social interaction (Fig. S6C), preference for novel animals in 3-chamber (Fig. S6D), or Y-maze (Fig. S6E).

Continuous homecage monitoring for 1 month revealed a non-significant ∼15% reduction (*P* = 0.2) in average locomotor activity (Fig. S6F), especially in the dark cycle, but no obvious change in daily locomotor activity rhythms in *Srrm2*^+/-^ mice (Fig. S6G). Activity on a free-spinning running wheel placed in the home cage was markedly reduced (∼3-fold) (Fig. 7A), while in the open field test, *Srrm2*^+/-^ mice showed ∼17% reduction in movement (total distance traveled; Fig. 7B, 7C), ∼40% reduction in jump counts (Fig. 7D), and ∼30% reduction in ratio of center/margin distance traveled (Fig. 7E). Effects observed in the open field test were notably stronger in female *Srrm2*^+/-^ mice compared to male mice (Table S6). In the forced swim test (FST), *Srrm2*^+/-^ mice showed ∼30% increase in immobility time (Fig. 7F). In the acoustic response assay, *Srrm2*^+/-^ mice showed reduced startle responses across a broad range of startle frequencies (Fig. 7G); however, there was little impairment in prepulse inhibition (Fig. 7H).

**Fig. 7:**
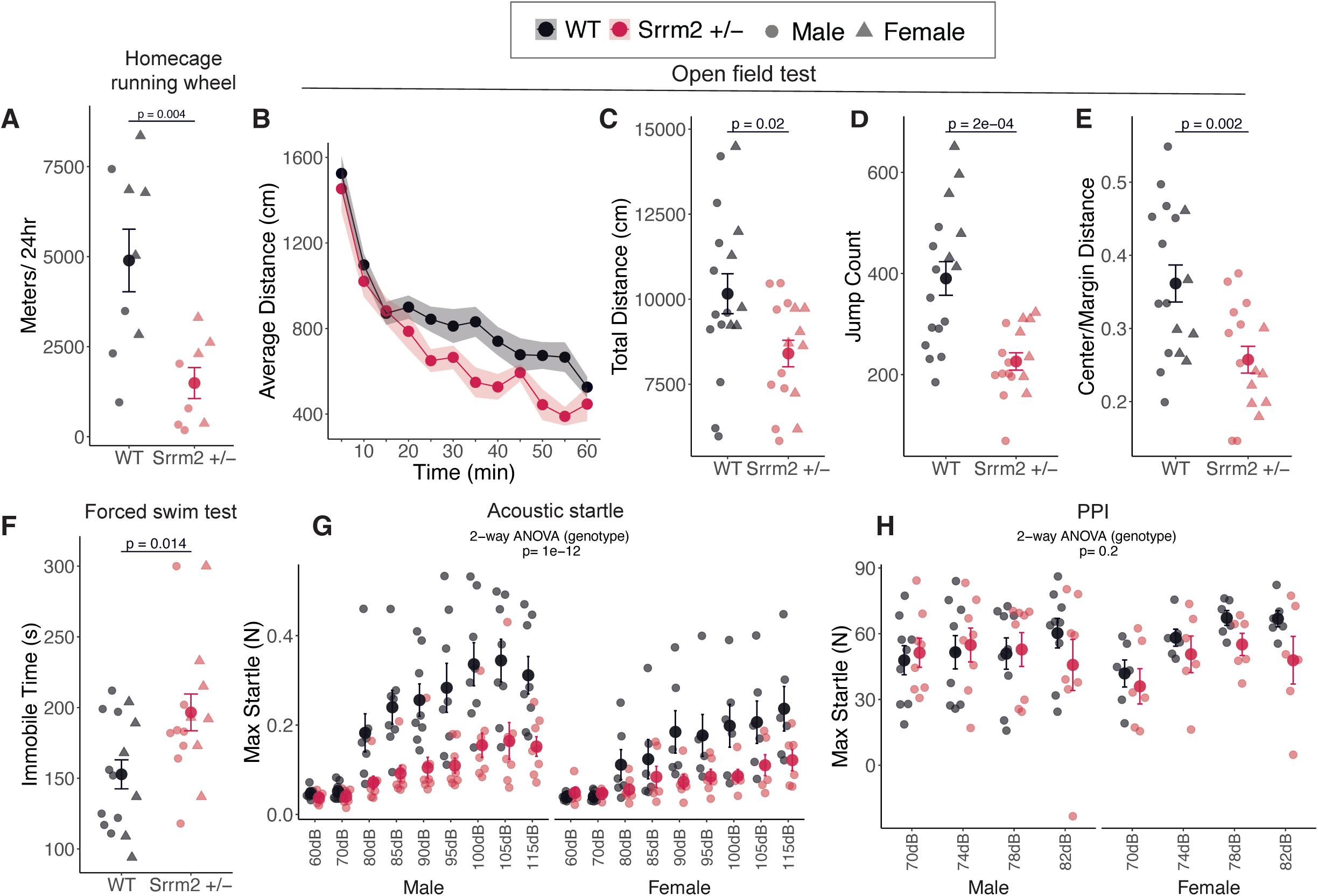
Behavioral abnormalities in *Srrm2*+/- mice. **(A)** Total running wheel distance of *Srrm2*+/- mice and WT littermate controls (n = 9 WT, n = 8 *Srrm2*+/-). Performance of *Srrm2*+/- mice in open field testing **(B–E)** (n = 17 WT, n = 16 *Srrm2*+/-); **(B)** distance traveled against time (averaged in 5-min bins), **(C)** total distance traveled summed over the duration of the assay, **(D)** total jump counts, and **(E)** ratio of total distance traveled in the center to that in the margin. **(F)** Immobility time (seconds) in the forced swim test (n = 15 WT, n = 15 *Srrm2*+/-). Auditory startle responses **(G–H)** (n = 14 WT, n = 14 *Srrm2*+/-); **(G)** startle force (newtons) and **(H)** percent inhibition of the auditory startle reflex with pre-pulse of the indicated dB. Statistical comparisons were performed using two-tailed Student’s t-tests for single-measure two-group comparisons and two-way ANOVA for multi-measure assays (acoustic startle and PPI). *P*-values are indicated in the panels. Error bars indicate SEM throughout.

Overall, *Srrm2*^+/-^ mice exhibit reduced locomotor activity and abnormal performance in behavioral assays commonly used to assess anxiety-like (center/margin distance) and depression-like behaviors (forced swim test, FST). It is possible that the observed behavioral impairments—including hypolocomotion, increased immobility in the FST, and reduced acoustic startle responses (measured as force exerted in Newtons by the mouse in the analysis chamber)—may reflect hypotonia (loss of muscle tone), a phenotype also reported in humans with *SRRM2*- ^5^ and *SYNGAP1-*haploinsufficiency ^26^. In this context, it is noteworthy SynGAP-γ is the only SynGAP isoform highly expressed in skeletal muscle.

### EEG changes reminiscent of humans with SCZ in *Srrm2*^+/-^ mice

Sleep disturbances and EEG abnormalities are common in humans with SCZ ^43,44^, and have been observed in genetic mouse models of SCZ ^45^. We therefore carried out 24-hour EEG recordings of *Srrm2*^+/-^ mice and WT littermate controls at 3- and 6-months of age.

Automated sleep scoring ^46^ during 12-hour light-dark cycles showed no significant difference in time spent in NREM sleep, REM sleep, or wake between *Srrm2*^+/-^ mutants and WT controls at 3 or 6 months of age (Fig. S7A).

Analysis of power spectral density during NREM sleep showed a widespread albeit non-significant attenuation in total EEG power (i.e. absolute power measured across all major frequency bands of brain oscillation) in S*rrm2*^+/-^ mice (Fig. 8A, Fig. S7C). Power in the sigma band (10-15 Hz) was particularly reduced in both parietal (p=0.053) and frontal (p=0.026) electrodes at 3-months of age (Fig. 8B, Fig. S7B).

**Fig. 8:**
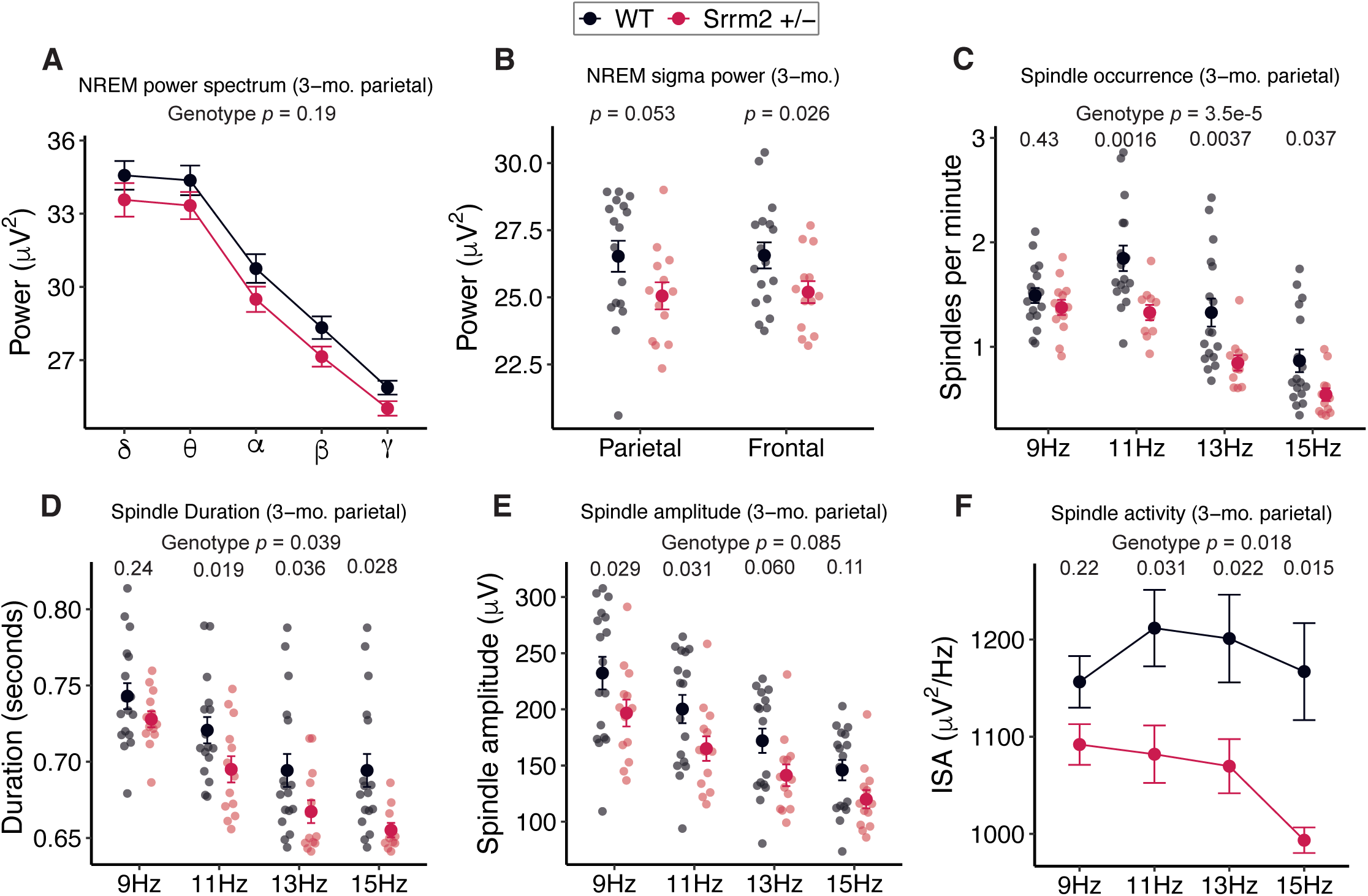
EEG changes reminiscent of humans with SCZ in *Srrm2*+/- mice. EEG recordings were performed in n = 13 WT and n = 17 *Srrm2*+/- mice. Signals were recorded for 24 hours beginning at ZT12, digitized at 1000 Hz, and bandpass filtered (Methods). **(A)** Absolute power of indicated frequency bands during NREM sleep in electrodes implanted in the parietal cortex of 3-month-old *Srrm2*+/- mice, compared to WT littermate controls. **(B)** Total sigma power in parietal and frontal cortex during NREM sleep at 3 months of age in *Srrm2*+/- mice and WT littermate controls. Spindle parameters in parietal cortex of 3-month-old *Srrm2*+/- mice compared to WT littermates: **(C)** sleep spindle occurrences (spindles per minute), **(D)** spindle duration, **(E)** spindle amplitude, and **(F)** integrated spindle activity. Spindles were detected using the Luna toolbox and segregated by center frequencies (9, 11, 13, and 15 Hz). EEG power density and spindle parameters were analyzed using linear mixed-effects models including genotype and frequency as fixed effects and subject ID as a random effect. The statistical impact of genotype was determined using likelihood ratio tests comparing full (genotype × frequency + subject ID) and reduced (frequency + subject ID) models. Statistical significance was defined as *P* < 0.05. Error bars indicate SEM throughout.

The decrease in sigma band power in *Srrm2*^+/-^ mice suggested reduced activity of sleep spindles, which are short bursts of 10-16 Hz oscillatory brain activity generated in the thalamus during NREM sleep ^43^. By analyzing sleep spindle events, we found that spindle occurrences (number of spindles per minute) (Fig. 8C, Fig. S7D), spindle duration (Fig. 8D, Fig. S7E), and spindle amplitude (Fig. 8E, Fig. S7F) were all substantially reduced in *Srrm2*^+/-^ mice, especially in the parietal electrode at 3 months of age (Fig. 8C–E). Correspondingly, integrated spindle activity (ISA, a metric that combines both duration and amplitude) was strongly reduced at the parietal electrode of 3-month-old *Srrm2*^+/-^ mice (Fig. 8F).

Sleep spindle occurrences, amplitude, and duration have consistently been found to be reduced (on average) in humans with SCZ ^43,44^; *Srrm2*^+/-^ mice therefore phenocopy a core EEG abnormality observed in SCZ. Because intact spindles correlate with cognitive function ^43,44^; impaired spindle activity may contribute to the cognitive impairment observed in humans with *SRRM2* LoF.

## DISCUSSION

*Srrm2*^+/-^ mice—an animal model of SCZ based on human genetics—exhibit several molecular and neurobiological phenotypes that overlap with abnormalities reported in humans with SCZ. These include downregulation of molecular pathways related to synapses, dysregulated dopamine signaling in striatum and PFC, deficits in ODCs and myelination, and reduced sleep spindles. In addition, our transcriptomic analysis reveals regional differences in putative brain activity, with hypoactivity in the PFC and hyperactivity in the hippocampus and somatosensory cortex (SSC). This pattern aligns with neuroimaging findings in individuals with SCZ ^47,48^, and broadly parallels the regional alterations in brain activity reported in *Grin2a*^+/-^ mice^49^.

Alternative splicing of mRNA is unusually prevalent in the brain ^50^ and is particularly affected in SCZ ^14^. It is therefore interesting that *SRRM2*, encoding an RNA splicing factor, has emerged as a large-effect LoF risk gene for SCZ ^2,3,51^. Mutations in and mislocalization of other nuclear RNA binding proteins (ex. TDP43, FUS) are linked to frontotemporal dementia ^52^, which can present with psychiatric symptoms including psychosis ^53,54^. Cytoplasmic mislocalization of SRRM2 itself—associated with disruption in speckles and splicing—has been observed in the brains of humans with frontotemporal dementia and Alzheimer’s disease ^33,55^, suggesting that disruptions in regulation of RNA splicing may contribute mechanistically to SCZ and multiple neurodegenerative diseases. In this context, it is notable that (i) several nuclear RNA binding proteins are elevated in synapse fractions of both *Srrm2*^+/-^ (Fig. 2C, Fig. S2C) and *Grin2a*^+/-^mouse models of SCZ (including Srrm2 itself in *Grin2a*^+/-^ synapses) ^49^, and (ii) chronic SCZ may have neurodegenerative and synapse loss components, especially later in life ^56^.

A major and unexpected conclusion stemming from our multi-omics study is that *Srrm2*^+/-^brains show reduced oligodendrocyte proportions and reduced levels of myelin proteins, particularly in the striatum. Dysmyelination is an emerging theme in the pathology of SCZ ^57^ and other CNS disorders ^58,59^. Neuroimaging ^60^, histopathological ^37^, transcriptomic ^14^, and genomic ^61^ studies have shown noted deficits in (i) ODC numbers, (ii) ODC- and myelin-related gene and/or protein expression, and (iii) myelination in the brains of humans with SCZ. In the *Srrm2*^+/-^mouse model we found reduction of ODC proportions and of myelin-related gene and protein expression in multiple regions of the brain, especially at 1 month of age. Thus, *Srrm2* heterozygous LoF mice phenocopy several cellular and molecular features of ODC depletion and reduced myelination seen in individuals with SCZ, raising the possibility that therapeutic strategies to promote myelination may be beneficial in some subsets of SCZ. In this context, we note that (i) in PheWAS of UK biobank exomes, *SRRM2* LoF is most strongly associated with myelination compared to all other traits ^62^, and (ii) *Srrm2* modulates early cell fate decisions ^34^, suggesting that oligodendrocyte deficits may originate from developmental shifts in precursor populations.

Dysregulated dopamine signaling in the striatum is one of the prevailing hypotheses for SCZ pathophysiology ^23^. We find considerable evidence of abnormal dopamine signaling and/or circuitry in the striatum of *Srrm2*^+/-^ mice, including elevated expression of dopamine signaling-related genes, and increased proportion of D1 and D2 SPNs in the striatum. In snRNA-seq measurements, D1 and D2 SPNs show downregulation of activity-regulated and dopamine-induced genes, implying reduced activity in these inhibitory neurons on average. Superficially, this does not align with a hyperdopaminergic state, which has been inferred in individuals with SCZ ^38,63^ and in the *Grin2a*^+/-^ SCZ mouse model ^49^. Further neurophysiological investigation of the striatum is needed to decipher the dopaminergic circuit dysfunction in *Srrm2* mutant mice.

Studies of both common and rare genetic variants ^2,22^, as well as postmortem brain tissue ^17,64^, have consistently pointed to synaptic dysregulation as a central pathomechanism in SCZ. We find that *Srrm2* haploinsufficiency exerts profound effects on synapses. By bulk RNA-seq, synapse-related gene sets are downregulated in all brain regions examined except for SSC, where they are upregulated, implying region-specific vulnerability, as previously observed in other genetic SCZ models ^49,65^. By synaptic proteomics, many key postsynaptic density (PSD) proteins are reduced in synapse fractions of adult *Srrm2*^+/-^ mice—most prominently, the γ isoform of SynGAP.

In addition to regulating Ras/Rap signaling, SynGAP is thought to play a structural role at the PSD, where it is highly concentrated (estimated local concentration of ∼100 µM) ^66,67^. It is therefore possible that the reductions of PSD proteins like Anks1b, Iqsec1, and Iqsec2—known SynGAP binding partners ^25,68^—reflect loss of scaffold support conferred by SynGAP-γ ^67,69^.

Heterozygous LoF mutations in *Syngap1* cause a neurodevelopmental syndrome characterized by intellectual disability, developmental delay, autism, and hypotonia—phenotypes that overlap with those observed in individuals with *SRRM2* haploinsufficiency ^5,26^. However, *Srrm2*^+/-^ mice exhibit reduced broadband EEG power and decreased locomotion, in contrast to the elevated EEG power and increased locomotion observed in global *Syngap1* haploinsufficiency ^70,71^. Although SynGAP isoforms are known to have distinct ^30,31^ and sometimes opposing ^72^ functions, this lack of concordance suggests that molecular changes beyond SynGAP-γ reduction, as well as broader circuit-level changes, contribute to the behavioral and network abnormalities in *Srrm2*^+/-^ mice.

Another prominent SynGAP-associated alteration in *Srrm2*^+/-^ brains is the marked elevation of full-length Agap3, a SynGAP-binding partner ^27–29^. Agap3 has been shown to participate in NMDA receptor-associated signaling and to modulate downstream Ras/ERK activation, in addition to acting as a GAP for Arf-family GTPases ^27^. Both SynGAP and Agap3 signaling converge on AMPAR trafficking and the control of synaptic strength ^73^. Prior work has shown that Agap3 restrains basal AMPAR surface expression, consistent with a role in limiting excitatory synaptic transmission ^27^. We therefore speculate that altered stoichiometry of the SynGAP/Agap3 complex shifts the balance of PSD-localized small GTPase signaling in a manner that reduces AMPAR surface abundance and influences excitatory synaptic transmission. Further investigations are needed to determine how these molecular and cellular mechanisms contribute to the systems and behavioral phenotypes of *Srrm2* mutant mice.

### Limitations of the study

All molecular, histological, and EEG studies were conducted exclusively in male mice, leaving potential sex-specific effects of *Srrm2* haploinsufficiency unresolved. Electrophysiological, spine density, and ultrastructural assessments are necessary to determine whether the synaptic and myelin protein alterations observed in *Srrm2*^+/-^ brains result in functional or structural deficits. Additional targeted validation of selected splicing events by RT-PCR/qPCR would further strengthen the transcriptomic findings.

## RESOURCE AVAILABILITY

### Lead contact

Further information and requests for resources and reagents should be directed to and will be fulfilled by the lead contact, Sameer Aryal.

### Materials availability

This study did not generate new reagents.

### Data and code availability

The original mass spectra and the protein sequence database used for searches have are available in the public proteomics repository MassIVE (http://massive.ucsd.edu) and with the accession MSV000095963. Raw bulk and single-nucleus RNA sequencing fastqs are available at NCBI GEO with the accession number GSE299937. Supplementary tables are available at https://doi.org/10.5281/zenodo.13910034

This paper does not report original code.

Any additional information required to reanalyze the data reported in this paper is available from the lead contact upon request.

## ACKNOWLEDGMENTS

The study was funded by the Stanley Family Foundation. Isoform-specific SynGAP ^30^ and Agap3 ^27^ antibodies were a kind gift from Yoichi Araki and Richard Huganir (Johns Hopkins). We thank Kris Dickson (Broad Institute) for feedback on the manuscript.

## AUTHOR CONTRIBUTIONS

Conceptualization, S.A. and M.S.; Methodology, S.A., M.S., S.A.C., H.K., and J.Q.P.; Validation, M.A.Y.; Formal Analysis, S.A., C.G., M.A.Y., H.K., A.N., A.S.A., and Y.W.; Investigation, C.G., M.A.Y., M.J.K., Z.F., N.G., K.S.B., N.S., A.G., K.J.S., O.S., and B.J.S.; Writing – Original Draft, S.A. and M.S.; Writing – Review & Editing, S.A. and M.S.; Visualization, S.A., C.G., M.A.Y.; Supervision, S.A., M.S., J.Q.P., S.P.M., S.A.C., and H.K.; Funding Acquisition, M.S.

## DECLARATION OF GENERATIVE AI AND AI-ASSISTED TECHNOLOGIES

During the preparation of this work the authors used ChatGPT/Gemini for readability improvements. After using this tool/service, the authors reviewed and edited the content and take full responsibility for the content of the published article.

## SUPPLEMENTAL INFORMATION

**Document S1. Figures S1–S7.**

**Table S1. Bulk RNA-seq differential expression and gene set enrichment analyses across brain regions and ages, related to Figure 1 and Figure S1.** Contains the results of DESeq2 analysis testing for differentially expressed genes in *Srrm2*+/-mice across the eight profiled brain regions at 1 and 3 months of age. Column suffixes indicate brain region and age. Additional sheets include GSEA results examining molecular pathways altered in the bulk RNA-seq data.

**Table S2. Synapse proteomics differential expression and pathway enrichment analyses in *Srrm2*+/- cortex, related to Figure 2 and Figure S2**. Contains the results of quantitative synapse proteomics comparing WT and *Srrm2*+/- cortical synapse fractions at 1 and 4 months of age, including differential protein expression statistics. Additional sheets include pathway enrichment analyses of the synaptic proteome using SynGO and MSigDB gene sets.

**Table S3. Isoform-level differential expression, aggregated gene-level isoform statistics, and alternative splicing analyses, related to Figure 3 and Figure S3.** Contains transcript isoform differential expression results, gene-level statistics derived from aggregation of isoform-level P values, and Leafcutter-based differential splicing analyses across brain regions and ages, testing for differences between *Srrm2*+/- and WT mice. Column suffixes indicate brain region and age. Additional sheets include rMATS summary counts of increased and reduced alternative splicing events for each event class in *Srrm2*+/- brains, as well as gene-level aggregation and Leafcutter-based differential splicing analyses of published RNA-seq data from human *SRRM2*-deficient iPSC-derived neurons.

**Table S4. Single-nucleus RNA-seq cell composition analyses and RNAscope-based quantification of oligodendrocyte and SPN proportions, related to Figure 4 and Figure S4.** Contains statistical results for *Srrm2*+/- versus WT cell type proportion analyses from single-nucleus RNA-seq in prefrontal cortex and striatum, as well as in situ RNAscope quantification of oligodendrocyte and SPN proportions and DAPI-positive cell density.

**Table S5. Single-nucleus RNA-seq differential expression and gene set enrichment analyses across neuronal and glial cell types, related to Figure 6**. Contains differential gene expression results across neuronal and glial cell types identified by single-nucleus RNA-seq in *Srrm2*+/- mice. Column suffixes indicate cell type. Additional sheets include GSEA results summarizing molecular pathways altered across the analyzed cell types.

**Table S6. Sex-stratified open field behavioral metrics in *Srrm2*+/- mice, related to Figure 7**. Contains female and male group means, percent differences, and t-test p values for center/margin distance ratio, total distance traveled, and jump count in the open field assay.

## FIGURE TITLES AND LEGENDS

**Fig. S1:**
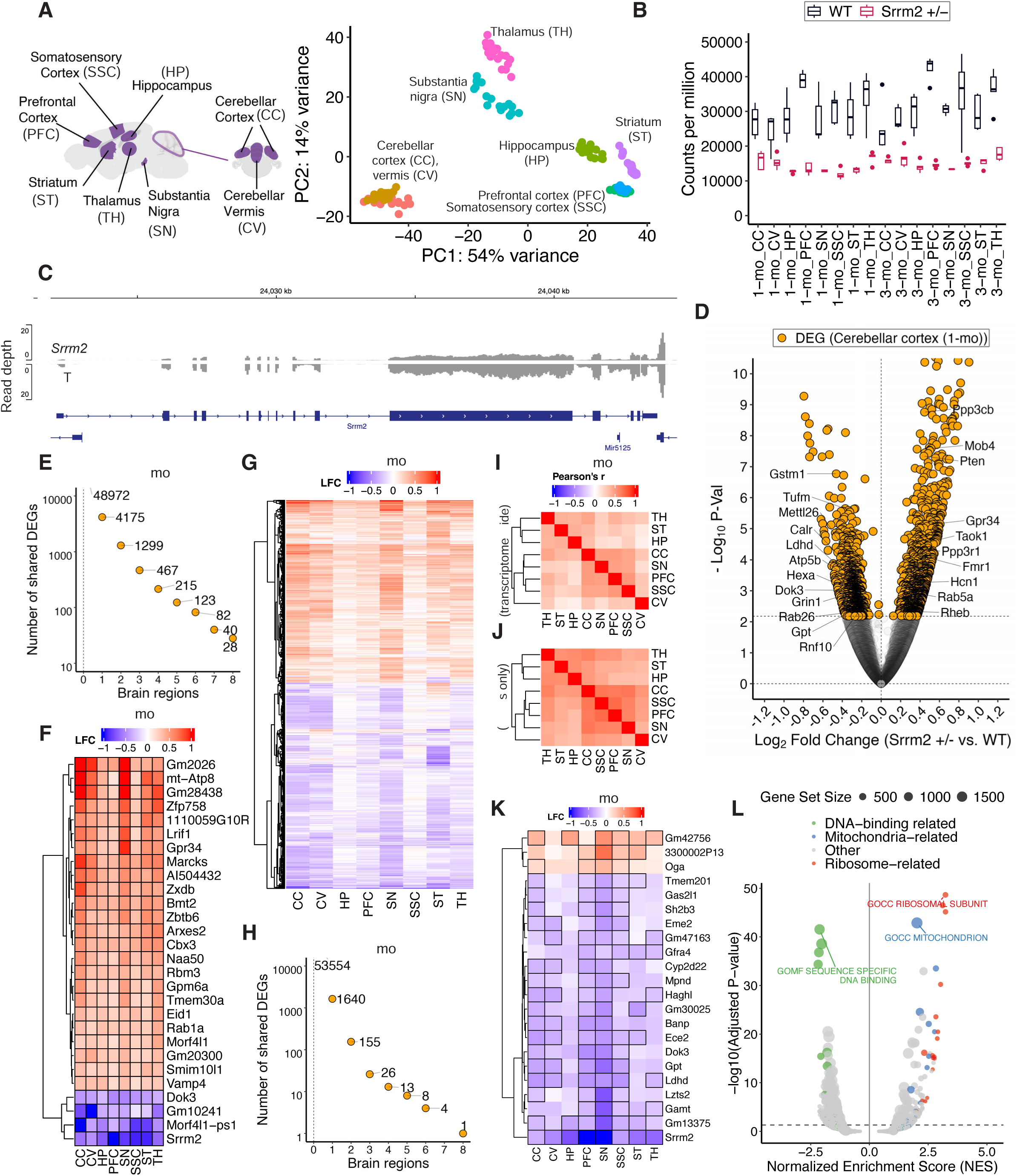
Large-scale and correlated transcriptomic changes across different brain regions in *Srrm2*+/- mice. RNA-seq and differential expression analyses were performed as described in Fig. 1 (5 WT and 5 *Srrm2*+/- per brain region per age; DESeq2 Wald test with Benjamini–Hochberg FDR correction). **(A)** The eight brain regions dissected (left) and principal component analysis (PCA) of RNA-seq datasets from those regions in 1- and 3-month-old *Srrm2*+/- mice and WT littermate controls (right). **(B)** Normalized counts (counts per million) of the *Srrm2* gene across all profiled brain regions in 1- and 3-month-old WT and *Srrm2*+/- mice. **(C)** Representative RNA-seq read density at the *Srrm2* locus in the PFC of 1-month-old *Srrm2*+/- mice and WT littermate controls, showing reduced read coverage across the gene body due to exon 9 deletion. **(D)** Volcano plot of gene expression changes in the cerebellar cortex of 1-month-old *Srrm2*+/-mice. Dotted horizontal line indicates FDR-adjusted *P* = 0.05. Genes shown in Fig. 1E are labeled. Number of shared DEGs across brain regions in 1-month-old **(E)** and 3-month-old **(H)** *Srrm2*+/-mice. The indicated numbers are not cumulative (e.g., genes DE in exactly seven regions do not include those DE in all eight regions, i.e. for 1-month-old *Srrm2*+/- mice, the 40 genes DE in exactly 7 brain regions does not include the 28 genes DE in all 8 regions, and the sum of 28, 40, 82, 123, 215 (=488) is the number of DEGs shared across at least four brain regions). Heatmap showing log2FCs of genes that are DE across all eight brain regions in 1-month-old *Srrm2*+/- mice **(F)**, and heatmap of log2FCs in the 1299 genes DE across exactly two brain regions in 1-month-old *Srrm2*+/- mice **(G)**. Pearson correlation of gene expression changes (log2FC) in 3-month-old *Srrm2*+/- mice **(I)** transcriptome-wide and **(J)** in the 1640 genes that are DE in any brain region. **(K)** Heatmap showing log2FCs in genes that are DE in four or more brain regions in 3-month-old *Srrm2*+/- mice. Only genes detected across all eight regions are included for hierarchical clustering. **L)** Volcano plot showing GSEA results for the *Age × Genotype* interaction across all profiled brain regions. GSEA was performed on preranked genes (ranked by the DESeq2 Wald test statistic); gene sets were considered significantly enriched at FDR-adjusted P < 0.05. Ribosome-related, mitochondria-related, and DNA-binding–related pathways are highlighted; the top changed gene set in each category is labeled.

**Fig. S2:**
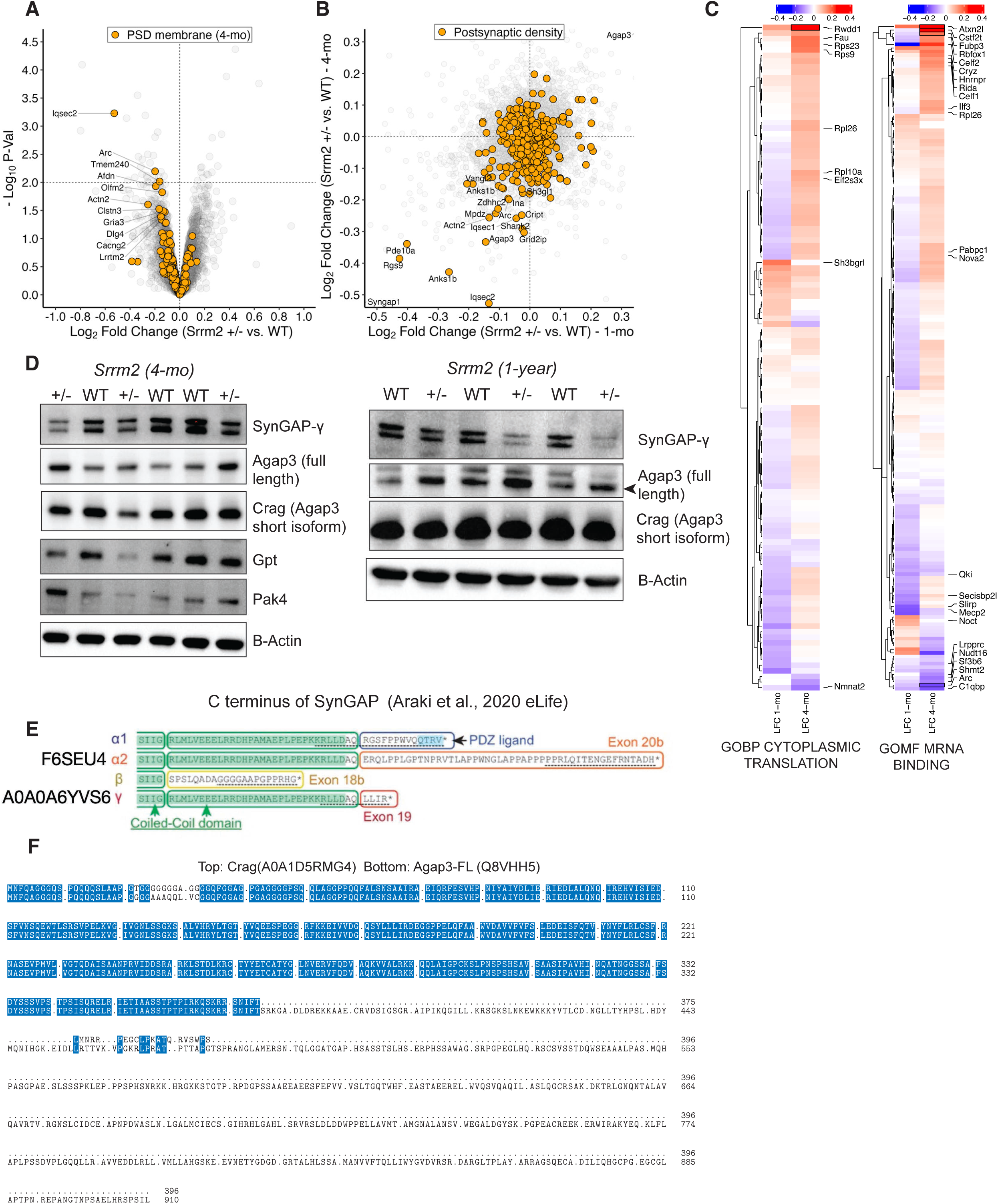
*Srrm2* LoF causes isoform-specific changes in SynGAP and Agap3 at the synapse. Synapse proteomics profiling and statistical analyses were performed as described in Fig. 2 (1-month: WT n = 4, *Srrm2*+/- n = 5; 4-month: WT n = 5, *Srrm2*+/- n = 4). **(A)** Volcano plot showing cortical synapse proteome changes in 4-month-old *Srrm2*+/- mice. Members of the SynGO “postsynaptic density membrane” gene set are highlighted. **(B)** Evaluation of log2FCs in 1-month-old *Srrm2*+/- synapses vs. those in 4-month-old synapses; individual components of the SynGO “postsynaptic density” gene set are highlighted. **(C)** Heatmap showing log2FCs in ribosomal proteins and RNA-binding proteins in 1- and 4-month-old *Srrm2*+/- mice synapses. Stroked cells indicate DEPs (*P* < 0.01, as defined in Fig. 2). Proteins with *P* < 0.1 in 1- or 4-month-old synapses (*Srrm2*+/- vs. WT) are labeled. **(D)** Western blots showing levels of various proteins of interest in synapse fractions from 4-month-old and 1-year-old *Srrm2*+/- mice and WT littermates. **(E)** Multiple sequence alignment of the C-terminus of the four Uniprot-annotated isoforms of the SynGAP protein. **(F)** Multiple sequence alignment of the protein isoforms of Agap3.

**Fig. S3:**
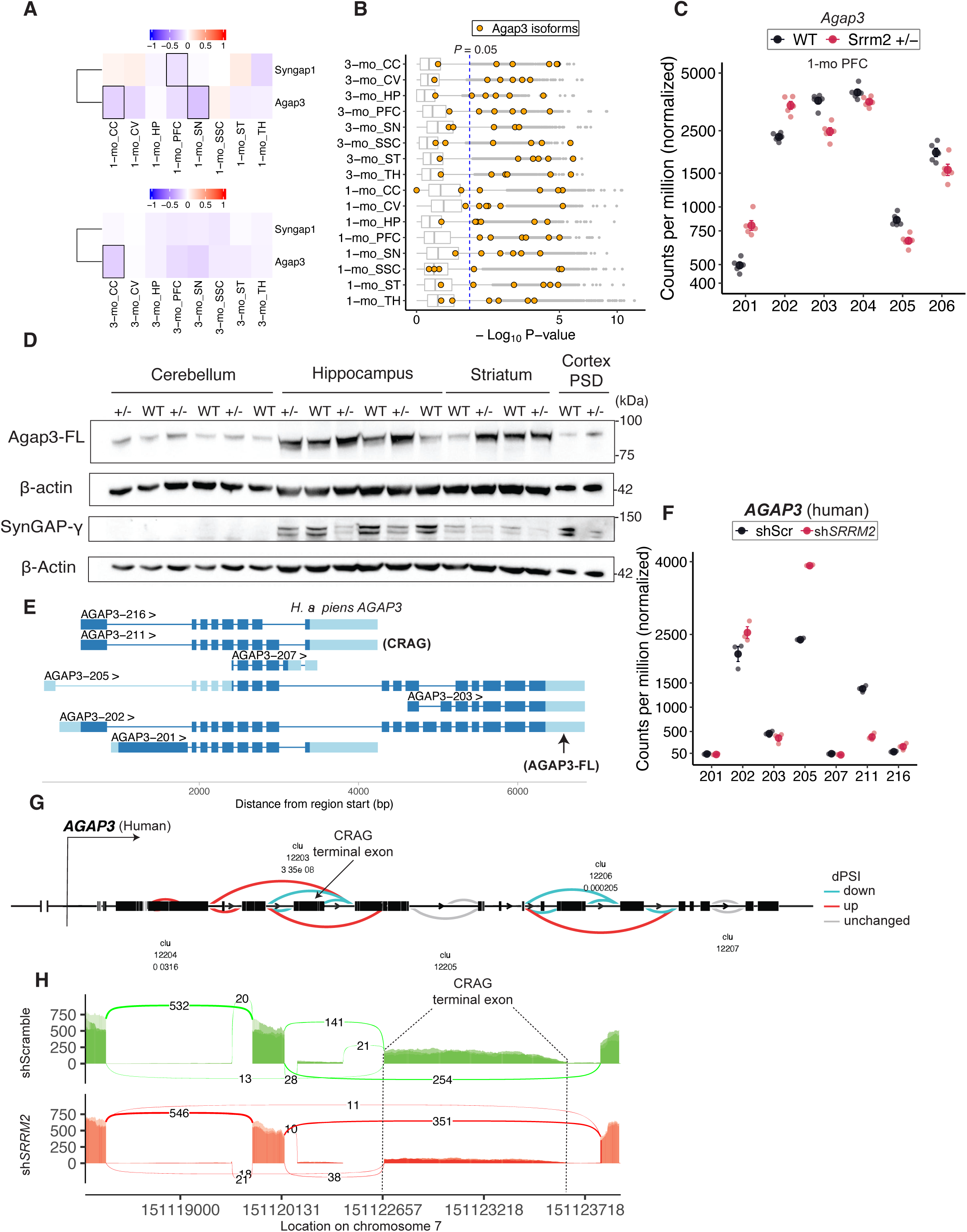
Altered Agap3 mRNA splicing across the brain of *Srrm2*+/- mice. Isoform-level differential expression and *P*-value aggregation were performed as described in Fig. 3 (Kallisto pseudoalignment, Sleuth isoform DE, Lancaster method for gene-level aggregation). Differential splicing analysis was performed using Leafcutter with likelihood ratio tests and FDR correction. **(A)** Gene expression changes (log2FC) in *Syngap1* and *Agap3* across brain regions; stroked cells indicate FDR-adjusted P < 0.05. **(B)** Nominal *P*-value distributions of the six mRNA isoforms of *Agap3* across brain regions in 1-and 3-month-old *Srrm2*+/- mice. **(C)** Counts of individual mRNA isoforms of *Agap3* in the PFC (representative brain region) of 1-month-old *Srrm2*+/- mice and WT littermates. **(D)** Western blot examining Agap3-FL and SynGAP-γ levels in P2 fractions of the cerebellum, hippocampus, and striatum of *Srrm2*+/- mice and WT littermate controls. The cortical synapse fraction (“cortex PSD”) serves as a positive control. **(E)** Gene models showing exonic and intronic regions of Ensembl-annotated H. sapiens *AGAP3* isoforms expressed in human iPSC-derived neurons. Isoforms with counts per million > 1 in RNA-Seq of human iPSC-derived neurons, obtained from Wu (2024) ^33^, are shown. **(F)** Counts of mRNA isoforms of *AGAP3* in human iPSC-derived neurons treated with scramble shRNA (shScr) or shRNA targeting *SRRM2* (shSRRM2). Isoforms encompassing the 3′ end of *AGAP3* (including *202*, encoding full-length AGAP3) are upregulated, while isoform *211* (encoding CRAG) is downregulated. **(G)** Gene model highlighting splicing changes in *AGAP3* identified by Leafcutter; “Clu” and associated dotted lines indicate intron clusters, and colors indicate changes in percent spliced in (PSI). A prominent splicing change (as indicated by low *P*-values) is found in the cluster encompassing the terminal exon of CRAG, which is skipped more frequently in human iPSC-derived neurons with *SRRM2* knockdown. **(H)** Sashimi plot showing altered exon usage in *AGAP3* following *SRRM2* knockdown in human iPSC-derived neurons. Numbers indicate reads spanning exon junctions. *SRRM2* knockdown causes an increase in reads that skip the terminal exon of CRAG (labeled), i.e. the number of reads aligning to the final exon of CRAG is reduced with *SRRM2* knockdown, while the number of reads spanning exon-junctions skipping this exon is elevated (254 in scramble-treated neurons vs. 351 in *SRRM2* knockdown neurons). Human iPSC data RNA-Seq data examining *SRRM2* knockdown (vs. scramble), which was not previously analyzed for effects on *AGAP3*, was obtained from Wu (2024) ^33^.

**Fig. S4:**
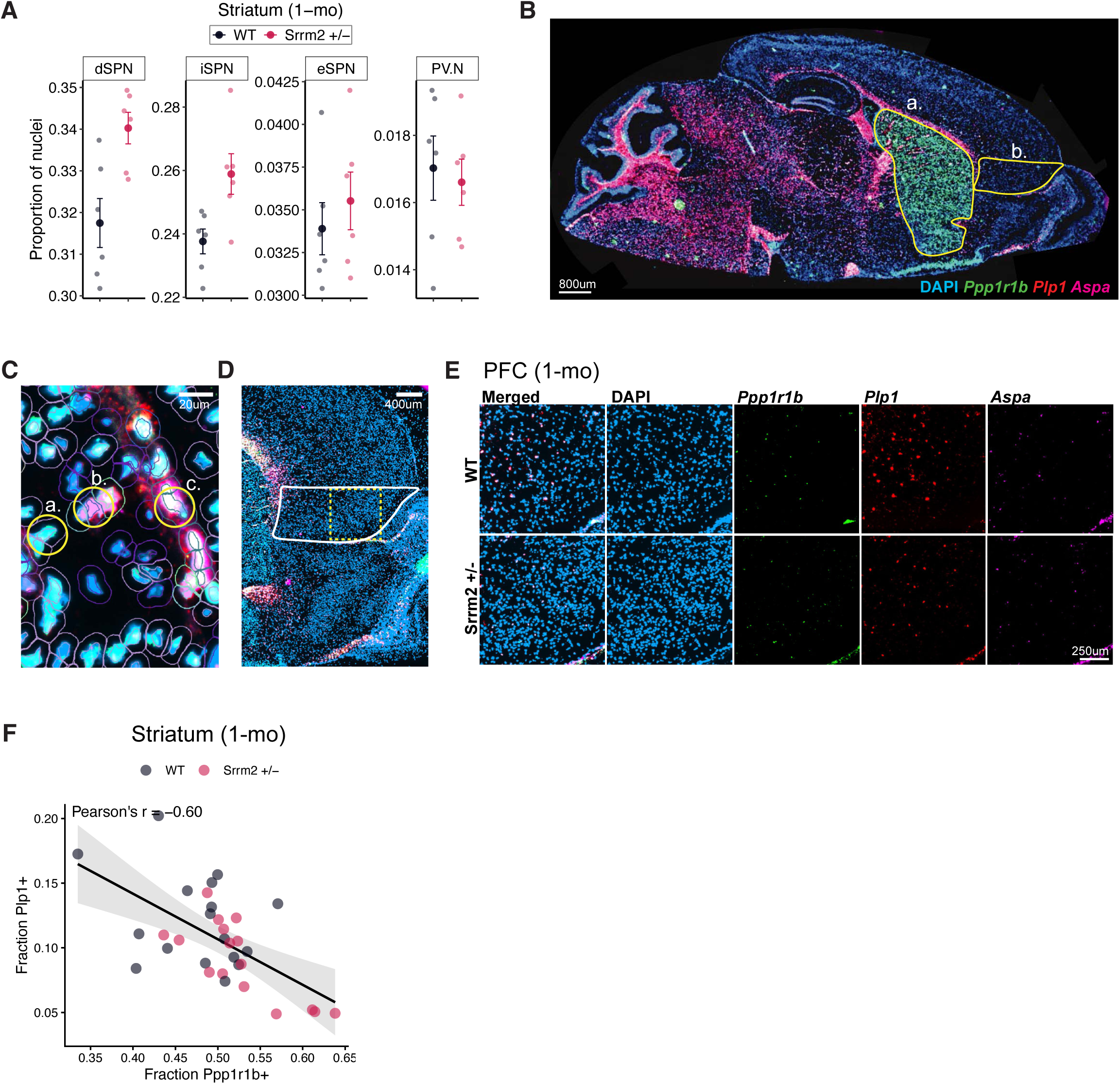
Decreased oligodendrocyte and increased SPN proportions in the *Srrm2*+/- brain. snRNA-seq cell composition analysis was performed as described in Fig. 4 (speckle propeller with arcsin normalization and Benjamini–Hochberg FDR correction; n = 6 WT and 6 *Srrm2*+/-per region). **(A)** Cell type proportion changes in inhibitory neuron subtypes of the striatum of 1-month-old *Srrm2*+/- mice. FDR-adjusted *P*-values testing for differences in proportions between WT and *Srrm2*+/- mice are indicated. dSPN = direct SPNs; iSPN = indirect SPNs; eSPN = eccentric SPNs; PV.N = parvalbumin neurons. Error bars indicate SEM. **(B)** Representative image of a full sagittal brain section used for RNAScope. Boxes denote (a) the striatum and (b) PFC regions of interest. **(C)** Representative cell detection and classification by QuPath. Inner circle indicates DAPI-defined nuclei and the outer circle estimates the cytoplasmic boundary. Examples show (a) *Ppp1r1b*□/*Aspa*□/*Plp1*□ (SPN), (b) *Ppp1r1b*□/*Aspa*□/*Plp1*□ (oligodendrocyte), and (c) *Ppp1r1b*□/*Plp1*□/*Aspa*□ (mature oligodendrocyte). **(D)** Representative sagittal section of the PFC. The white outline indicates the data collection area; the dotted yellow square indicates the high magnification region shown in (E). **(E)** Representative images of DAPI, *Ppp1r1b*, *Plp1*, and *Aspa* expression in WT vs. *Srrm2*+/-PFC. **(F)** Evaluation of proportion of *Ppp1r1b*□ (SPN) vs. *Plp1*□ (ODC) from RNAScope in *Srrm2*+/-and WT littermates. Data represent 6 WT and 6 *Srrm2*+/- animals at P29 (three sections per animal; each point = 1 section). Error bars indicate SEM.

**Fig. S5:**
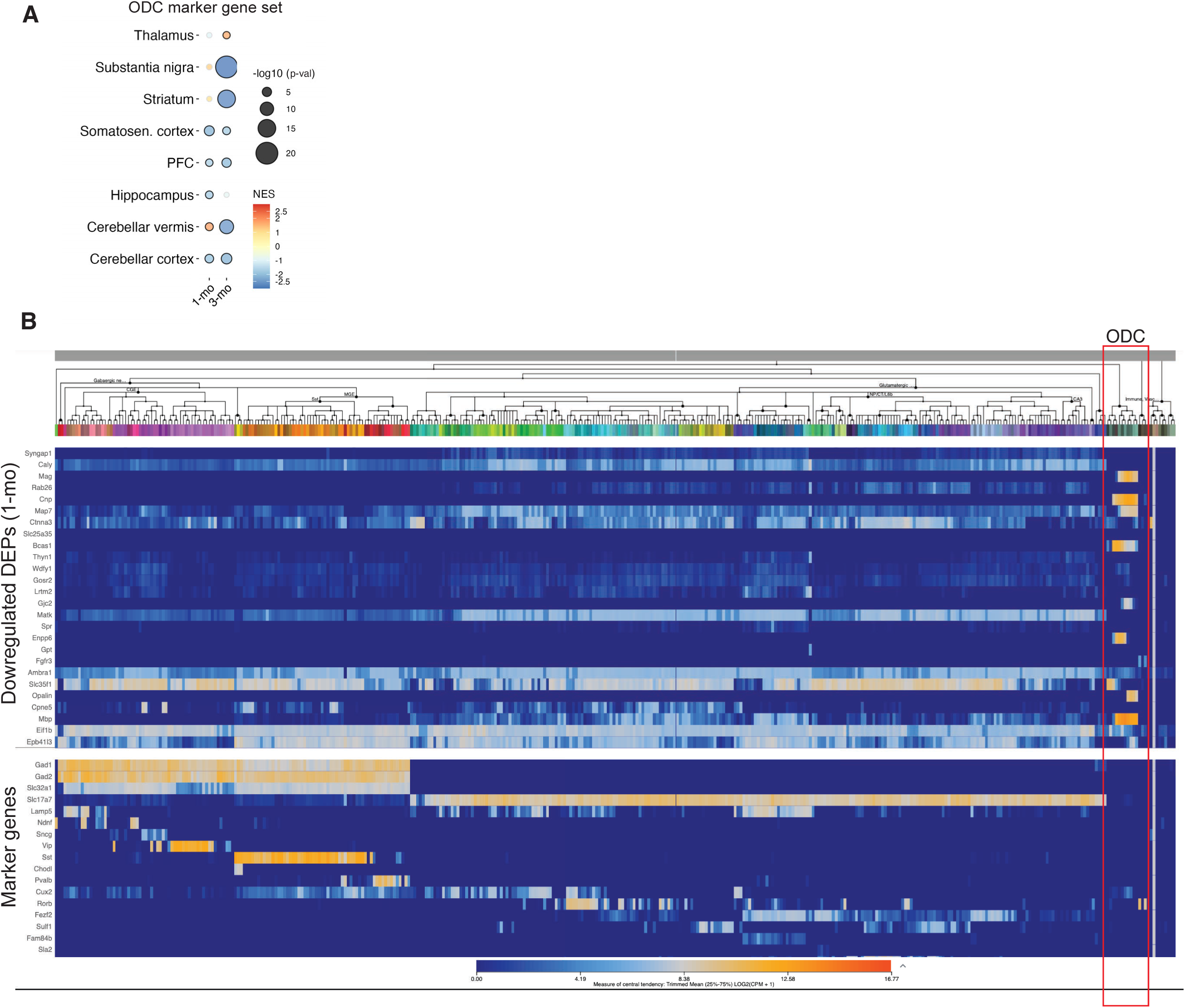
Decrease in myelin-related mRNAs and proteins in *Srrm2*+/- mice. Bulk RNA-seq and synapse proteomics datasets and statistical analyses were performed as described in Fig. 5. **(A)** Bubble plot showing bulk RNA-seq GSEA results for the “Lein_oligodendrocyte_marker” MSigDB gene set in 8 brain regions of 1- and 3-month-old *Srrm2*+/- animals. GSEA was performed on preranked genes (ranked by the DESeq2 Wald test statistic) using only the “Lein_oligodendrocyte_marker” gene set; Stroked circles indicate nominally significant enrichment (*P* < 0.05). **(B)** Evaluation of cell type–specific gene expression of the 27 reduced DEPs in 1-month-old *Srrm2*+/- synapses using the Allen Cell Types database. Of the 27 proteins, expression data were available for 26; 10 of 26 (e.g., Mag, Cnp, Map7) are highly enriched in oligodendrocytes (highlighted in red).

**Fig. S6:**
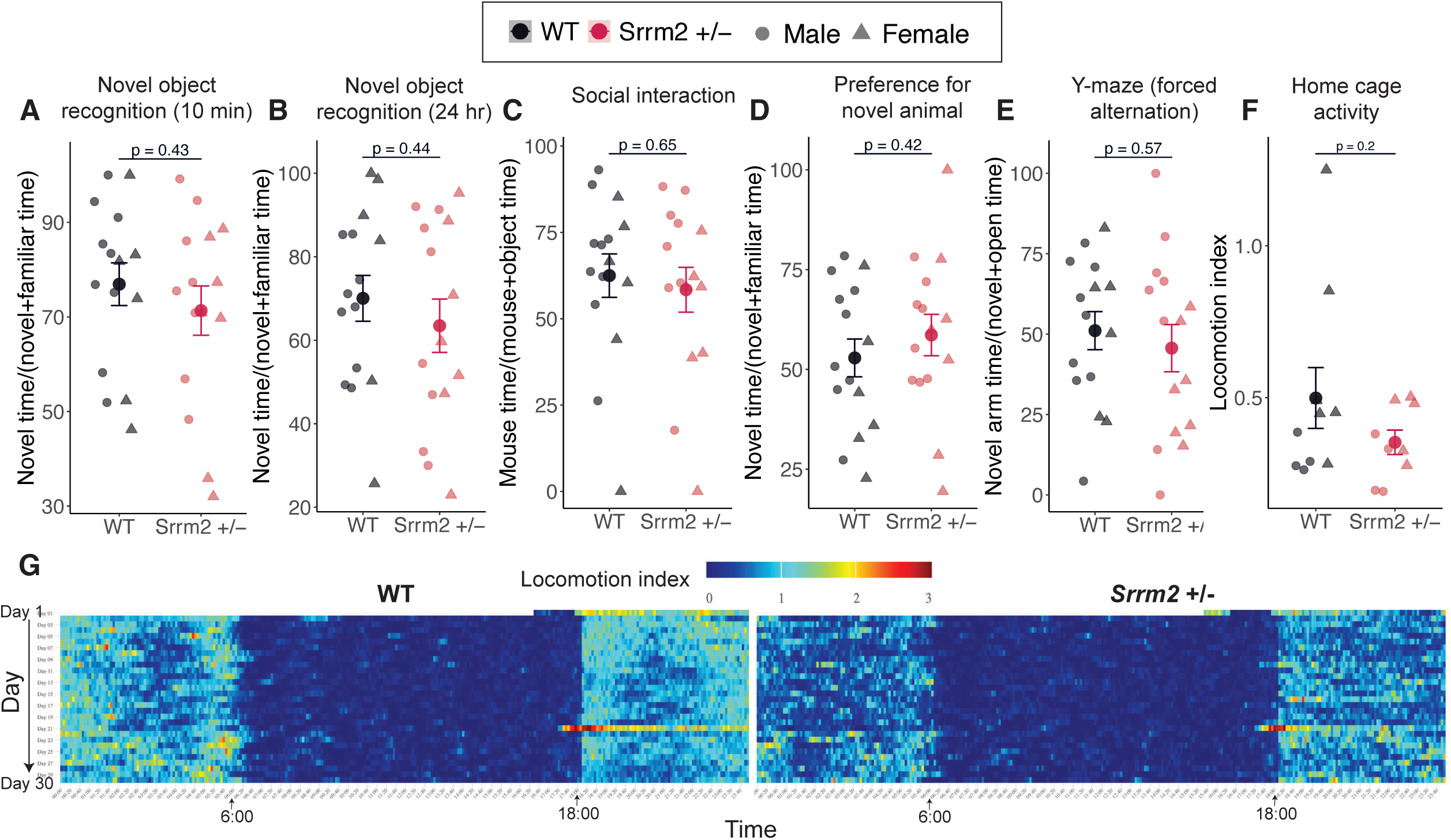
Behavioral abnormalities in *Srrm2*+/- mice. **(A–B)** Evaluation of novel object recognition in *Srrm2*+/- mice and WT littermate controls after **(A)** 10 minutes and **(B)** 24 hours of object exposure (n = 15 WT, n = 15 *Srrm2*+/-). **(C–D)** Evaluation of **(C)** three-chamber social interaction and **(D)** social recognition in *Srrm2*+/-mice compared to WT littermate controls (n = 15 WT, n = 15 *Srrm2*+/-). **(E)** Performance of *Srrm2*+/- mice in the Y-maze assay of spatial memory (n = 15 WT, n = 15 *Srrm2*+/-). Home cage activity measurements over 30 days in WT and *Srrm2*+/- mice (n = 9 WT, n = 9 *Srrm2*+/-): **(F)** average home cage activity across one month and **(G)** average activity across the duration of a day. Statistical comparisons were performed using two-tailed Student’s t-tests for single-measure two-group comparisons and two-way ANOVA for multi-measure assays where applicable. *P*-values testing for genotype differences are shown in the panels. Error bars indicate SEM throughout.

**Fig. S7:**
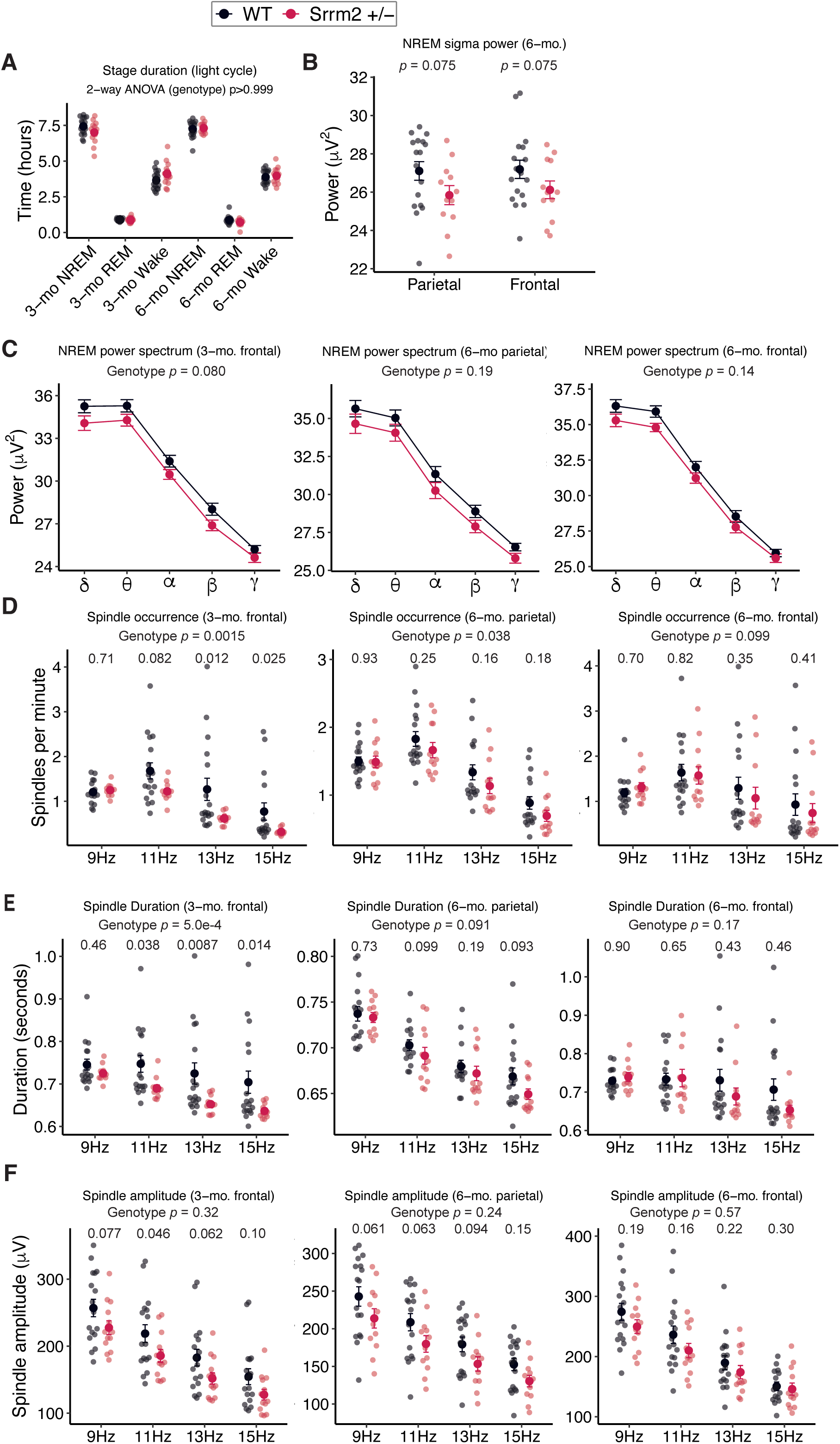
EEG changes reminiscent of humans with SCZ in *Srrm2*+/- mice. EEG recordings and statistical analyses were performed as described in Fig. 8 (n = 13 WT, n = 17 *Srrm2*+/-). **(B)** Time spent in REM sleep, NREM sleep, and wake (hours) by *Srrm2*+/- mice and WT littermate controls. Statistical significance was assessed using two-way ANOVA; F-test *P*-values testing for genotype effects are shown. **(C)** Total sigma power in parietal and frontal cortex during NREM sleep at ∼6 months of age in *Srrm2*+/- mice and WT littermate controls. **(D)** Absolute power of indicated frequency bands during NREM sleep in electrodes implanted in the indicated brain region at the indicated age in *Srrm2*+/- mice compared to WT littermates. Sleep spindle parameters in the indicated brain region at ∼3 and ∼6 months of age: **(D)** spindle occurrences (spindles per minute), **(E)** spindle duration, and **(F)** spindle amplitude. Spindles were segregated by center frequencies. EEG power density and spindle parameters were analyzed using linear mixed-effects models including genotype and frequency as fixed effects and subject ID as a random effect, as described in Fig. 8. Statistical significance was defined as *P* < 0.05. Error bars indicate SEM throughout.

## STAR METHODS

### KEY RESOURCES TABLE

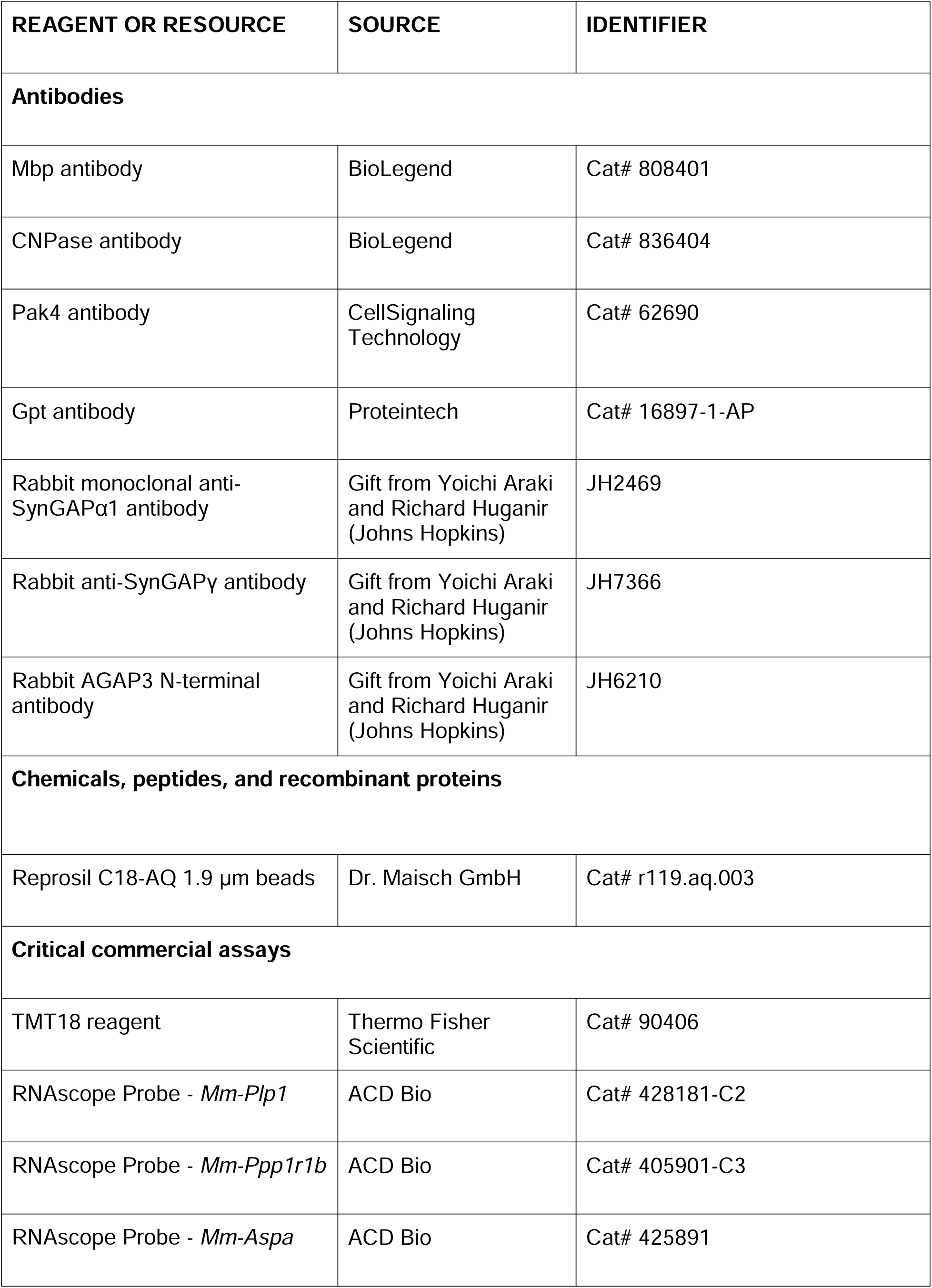
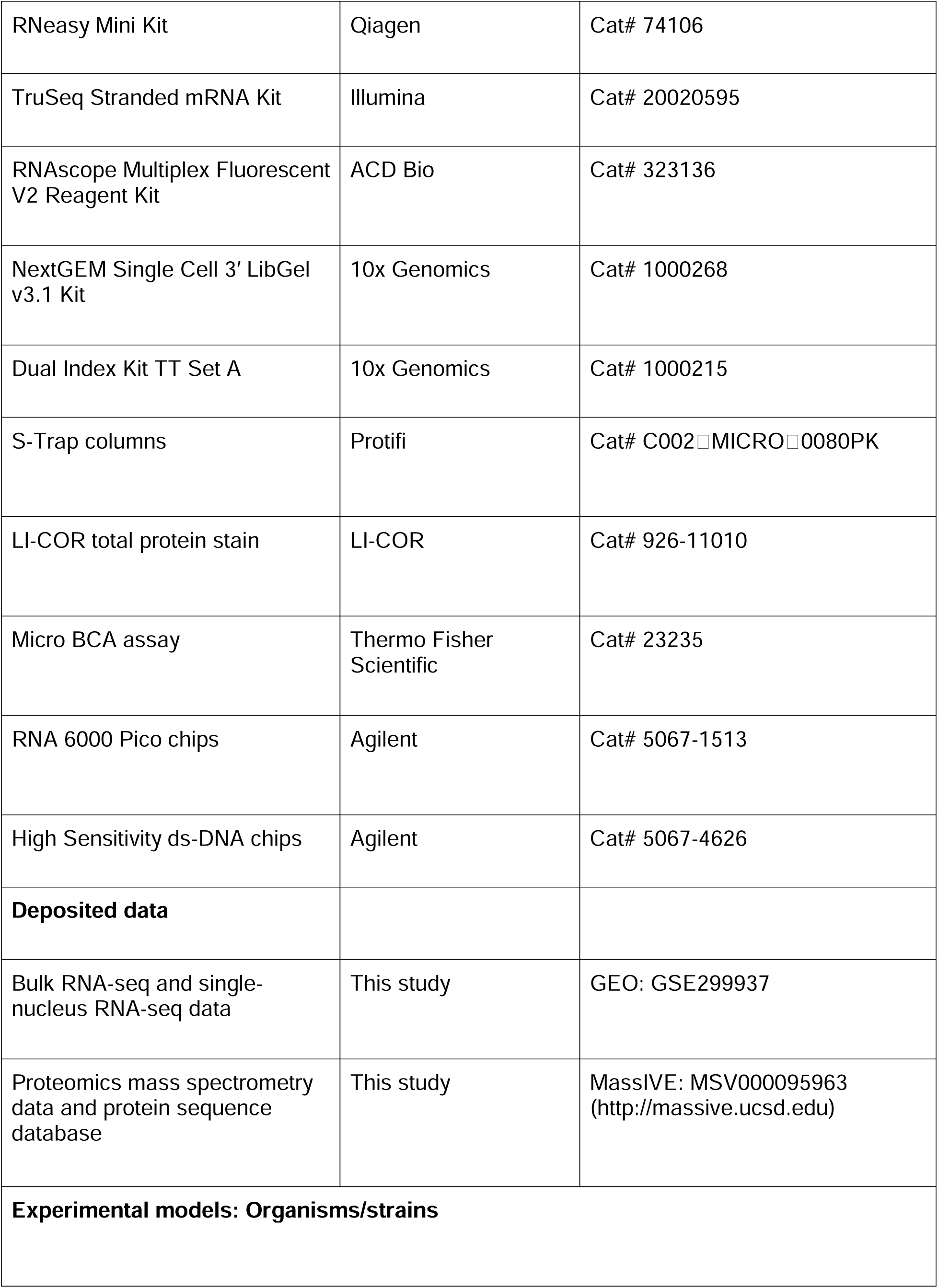
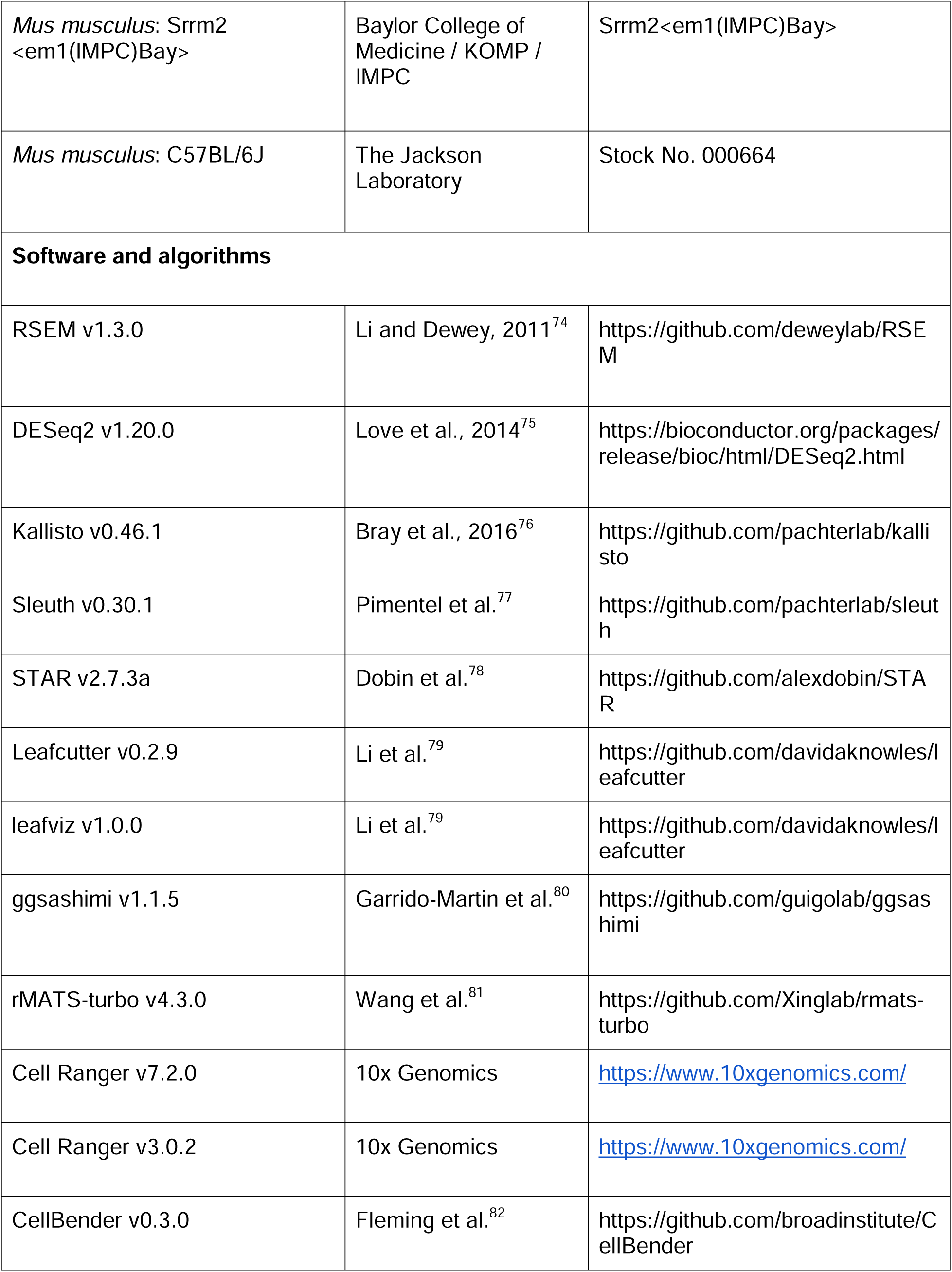
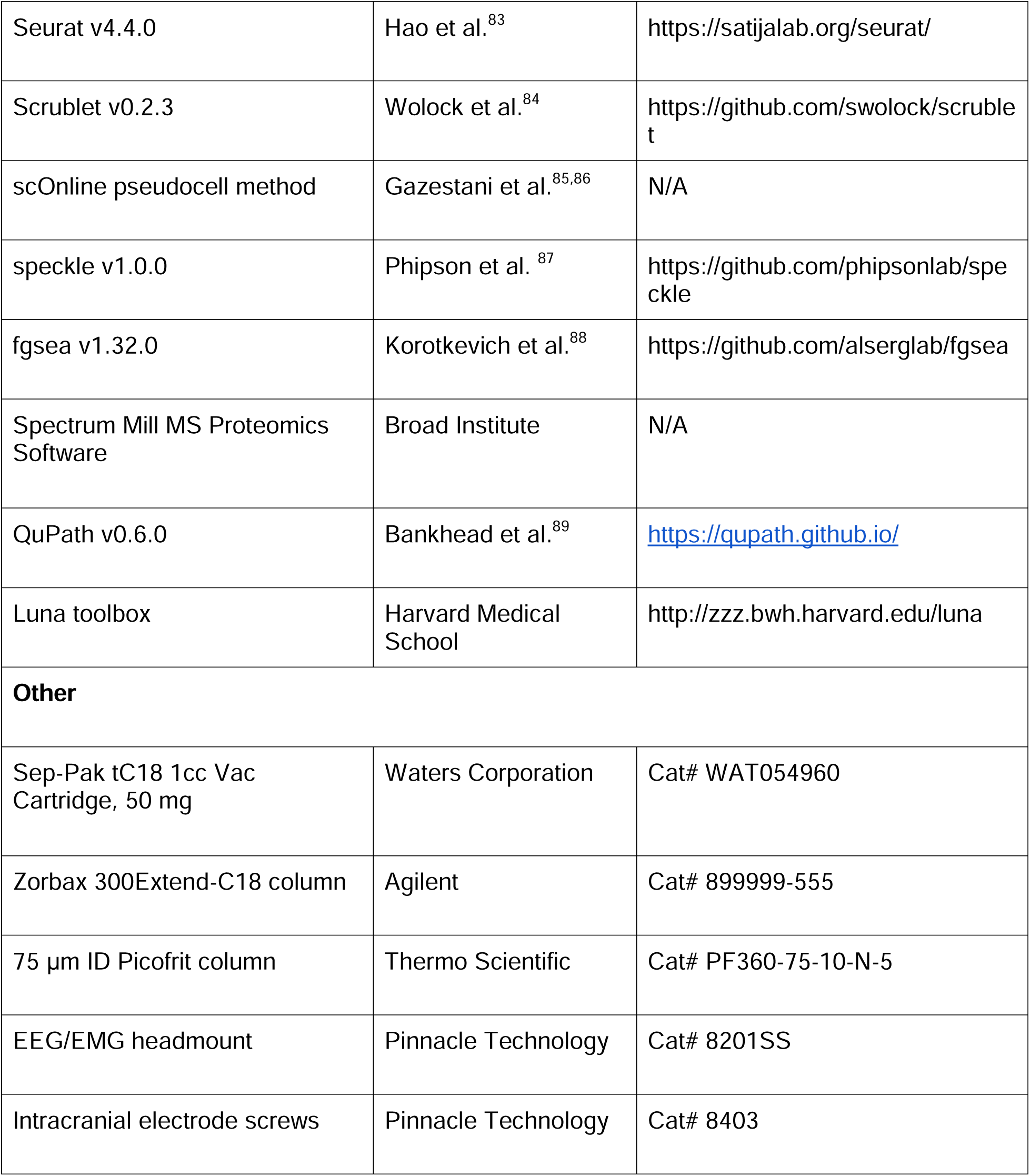

### EXPERIMENTAL MODEL AND STUDY PARTICIPANT DETAILS

#### Mouse line generation, breeding strategy, and ethical considerations

We studied the previously generated *Srrm2 ^em1(IMPC)Bay^*mutant that harbors a deletion of exon 9 (introduced by the Cas9-RNA guided nuclease) in the *Srrm2* locus ^90^, which we confirmed by Sanger sequencing and PCR. This line was generated by the knockout mouse program (KOMP) / International Mouse Phenotyping Consortium (IMPC) at Baylor College of Medicine (Houston, TX) ^90^. While heterozygous mutants of this mouse line are generally healthy, homozygous knockouts show complete penetrance of preweaning lethality ^90^; so we focused our investigations on the heterozygous (+/-) genotype, which is also the genotype linked to elevated risk of SCZ and the *SRRM2* LoF syndrome.

Cryopreserved germplasm (sperm samples) heterozygous for the *Srrm2 ^em1(IMPC)Bay^* allele were obtained from John Seavitt (Baylor College of Medicine) and mutant mice were re-derived by in-vitro fertilization at Harvard Genome Modification Facility (Cambridge, MA). Mutant mice were back-crossed with C57BL/6J (The Jackson Laboratory, #000664) for two generations, after which female mutant mice were bred with C57BL/6J males to generate experimental animals (*Srrm2*^+/-^ and WT littermates). To minimize genetic drift, breeders were periodically refreshed by pairing fresh ∼4-month-old *Srrm2 ^em1(IMPC)Bay^* mutant females with freshly obtained C57BL/6J males from the Jackson Laboratory.

Mice were housed in AAALAC-approved facilities on a 12-hour light/dark cycle, with food and water available *ad libitum*. All procedures involving mice were approved by the Broad Institute IACUC (Institutional Animal Care and Use Committee; protocol #0259-10-19-2) and conducted in accordance with the NIH Guide for the Care and Use of Laboratory Animals.

Male mice were used for biochemical and molecular studies (RNA-seq, proteomics, synapse fractionation, western blotting), as well as for EEG studies. Both male and female mice were used for behavioral studies.

### METHOD DETAILS

#### Brain perfusion and dissection

Mice were transcardially perfused under 3% isoflurane anesthesia with Hank’s Balanced Salt Solution (HBSS). The perfused brains were extracted, frozen in liquid nitrogen vapor, and stored at -80°C as described in detail in Vanderburg et al. ^91^. Regional microdissections were performed on a cryotome (Leica CM3050S) as described in Farsi et al. ^49^. Briefly, after cerebellum removal and midsagittal hemisection, the medial prefrontal cortex was hand-dissected from the exposed medial surface using a precooled microscalpel. Hemispheres were then remounted onto a new cryostat chuck with O.C.T. freezing medium, and 30 µm coronal sections were progressively removed to expose deeper structures. The dorsal hippocampus and somatosensory cortex were hand-dissected using a microscalpel, while the thalamus and dorsal striatum were isolated using a 0.15 cm diameter biopsy punch. All dissected tissue was stored at –80°C.

#### RNA isolation and bulk RNA-seq library preparation

Dissected brain tissue was lysed, and total RNA isolated using RNeasy Mini Kit (Qiagen) following the manufacturer’s instructions. RNA concentration was measured using Nanodrop, and RNA quality / RNA integrity number assessed using RNA 6000 Pico Electrophoresis chips on an Agilent Bioanalyzer. Isolated total RNA was stored at -80°C until bulk RNA-seq library preparation.

Bulk RNA-seq cDNA libraries were prepared using a TruSeq Stranded mRNA Kit (Illumina). The fragment sizes of the cDNA libraries were analyzed using High Sensitivity DNA chips (Agilent) on a 2100 Bioanalyzer Instrument (Agilent), and their concentrations measured using Qubit fluorometer. A 10 nano molar (nM) normalized library was pooled, and paired-end sequencing was performed on a NovaSeq S2 (Illumina) with 50 bases each for reads 1 and 2 and 8 bases each for index reads 1 and 2.

Bulk RNA-seq libraries were prepared from 5 WT and 5 *Srrm2*^+/-^ mice for each brain region and age. Three 3-month SN samples (1 WT and 2 *Srrm2*^+/-^) were excluded from sequencing due to failure in library preparation.

#### Biochemical preparations

##### Preparation of total homogenate extracts

Mice were euthanized using CO₂ inhalation, after which the cortex, hippocampus, and cerebellum were rapidly dissected, flash-frozen in liquid nitrogen, and stored at −80 °C until processing.

Frozen brain tissue was thawed on ice and Dounce-homogenized in ice-cold homogenization buffer (5 mM HEPES, pH 7.4; 1 mM MgCl□; 0.5 mM CaCl□; supplemented with protease and phosphatase inhibitors).

For total homogenate preparation, sodium dodecyl sulfate (SDS) was added to the homogenate to a final concentration of 2%. The sample was probe-sonicated, clarified by centrifugation at 15,000 g for 10 min at room temperature, and the supernatant was collected as the total protein extract.

##### Purification of P2 (crude synaptosomal) fraction

For preparation of the P2 fraction, SDS was not added to the homogenate. Instead, the homogenate was centrifuged at 1,400 g for 10 min at 4 °C to remove nuclei and large debris (P1 fraction). The resulting supernatant was subsequently centrifuged at 13,800 g for 10 min at 4°C. The resulting pellet (P2 fraction), corresponding to the crude synaptosomal fraction enriched in synaptosomes, mitochondria, and other heavy membrane-associated components, was resuspended in 0.32 M sucrose, 6 mM Tris-HCl (pH 7.5). The P2 fraction was either analyzed by SDS-PAGE or used for synaptic fractionation, as described below.

##### Preparation of synaptic (postsynaptic density) fractions

To obtain synaptic fractions, the P2 fraction was gently layered onto a discontinuous sucrose gradient (0.85 M, 1.0 M, and 1.2 M sucrose in 6 mM Tris-HCl, pH 8.0) and ultracentrifuged at 82,500 g for 2 h at 4 °C. The synaptosome fraction, which sediments at the 1.0 M / 1.2 M sucrose interface, was collected and diluted with an equal volume of ice-cold 1% Triton X-100 (in 6 mM Tris-HCl, pH 8.0). After incubation on ice for 15 min, samples were ultracentrifuged at 32,800 g for 20 min at 4 °C. The resulting pellet, corresponding to a postsynaptic density (PSD)–enriched fraction, was resuspended in 1% SDS. The purification of synapse fractions was validated by western blotting of synaptic markers, demonstrating ∼20-fold enrichment of PSD-95 relative to total homogenate. Although this fraction is often referred to as a “PSD fraction,” we refer to it throughout this manuscript as the “synapse fraction,” as it contains abundant PSD proteins as well as presynaptic active zone (cytomatrix of the active zone) proteins. This terminology is consistent with our prior published work using the same fractionation approach ^17,49,65^.

An aliquot was used to determine protein concentration using the micro BCA assay (Thermo Fisher Scientific), and the remaining material was stored at −80 °C prior to mass spectrometry or western blot analysis.

##### Processing of synaptic fractions for mass spectrometry

Eighteen synaptic fractions from WT and *Srrm2*^+/-^ mouse cortices were analyzed using tandem mass tag (TMT) isobaric labeling for quantification (1-mo WT n = 4, 1-mo *Srrm2*^+/-^ n = 5, 4-mo WT n = 5, 4-mo *Srrm2*^+/-^ n = 4).

The protein in synapse fraction samples (in 1% SDS) was reduced using 5 mM dithiothreitol and alkylated using 10 mM iodoacetamide at room temperature. The denatured, reduced, alkylated protein samples were then processed using S-Trap sample processing technology (Protifi) following manufacturer’s instructions. The proteins were bound to the S-Trap column via centrifugation and contaminants/detergents were washed away. Sequential digestion steps were then performed on column using 1:20 enzyme to substrate ratio of Lys-C for 2 hours and Trypsin overnight at room temperature.

Following digestion and desalting, 15 μg of each sample was labeled with TMT18 reagent; the TMT18 plex was constructed by randomly assigning the samples from each group to channels within the plex. After verifying successful labeling of more than 99% label incorporation, reactions were quenched using 5% hydroxylamine and pooled. The TMT18 labeled peptides were desalted on a 50 mg tC18 SepPak cartridge and fractionated by high pH reversed-phase chromatography on a 2.1 mm x 250 mm Zorbax 300 extend-c18 column (Agilent). One-minute fractions were collected during the entire elution and fractions were concatenated into 12 fractions for LC-MS/MS analysis.

One microgram of each proteome fraction was analyzed on a Exploris 480 QE mass spectrometer (Thermo Fisher Scientific) coupled to a Vanquish Neo LC system (Thermo Fisher Scientific). Samples were separated using 0.1% Formic acid / Acetonitrile as buffer A and 0.1% Formic acid / Acetonitrile as buffer B on a 27cm 75um ID picofrit column packed in-house with Reprosil C18-AQ 1.9 mm beads (Dr Maisch GmbH) with a 90 min gradient consisting of 1.8-5.4% B in 1 min, 5.4-27% B for 84 min, 27-54% B in 9 min, 54-81% B for 1 min followed by a hold at 81% B for 5 min. The MS method consisted of a full MS scan at 60,000 resolution and a normalized AGC target of 300% and maximum inject time of 10ms from 350-1800 m/z followed by MS2 scans collected at 45,000 resolution with a normalized AGC target of 30% with a maximum injection time of 105 ms and a dynamic exclusion of 15 seconds. Precursor fit filter was used with a threshold set to 50% and window of 1.2 m/z. The isolation window used for MS2 acquisition was 0.7 m/z and 20 most abundant precursor ions were fragmented with a normalized collision energy (NCE) of 34 optimized for TMT18 data collection.

##### Western blotting

Total extracts, P2 fractions, and synaptic fractions were prepared as described, and their protein concentrations measured using Bicinchoninic acid assay (BCA; Thermo Fisher). Laemmli buffer (6X; Boston Bioproducts BP-111R) was added to the samples at a final concentration of 1X, and the samples heated at 95°C for 5 minutes.

SDS-PAGE was carried out by electrophoresing the samples on a denaturing 4-12% NuPage gradient gels (Thermo Fisher). Gels were transferred to nitrocellulose membranes (Bio-Rad TransBlot Turbo), and total protein staining (LiCor 926-11010) was performed. Membranes were then blocked with 5% milk in Tris-buffered saline with 0.1 % Tween 20 (TBST). Membranes were incubated with primary antibodies overnight at 4C, washed three times with TBST, and then incubated with secondary HRP-conjugated antibodies for 1-2 hours at room temperature. After additional washing (4 × 5 minutes), membranes were imaged on ChemiDoc XRS+ (Bio-Rad) and analyzed using Image Lab Software (Bio-Rad).

The antibodies used were as follows: Mbp (Biolegend 808401), CNPase (Biolegend 836404), Pak4 (Cell Signaling 62690), Gpt (ProteinTech 16897-1-AP). Isoform-specific SynGAP 30 and Agap3 27 antibodies were a kind gift from Yoichi Araki (Johns Hopkins).

##### Nuclei purification and snRNA-seq library preparation

Nuclei were purified by dissociating dissected brain tissue in a 1% Triton-X based buffer, followed by FACS enrichment, as detailed in ^92^. Purified nuclei were loaded on a 10x Chromium v3.1 system (10x Genomics) and library preparation (Single Cell 3□; 10X Genomics) was performed according to the manufacturer’s protocol. cDNA fragment sizes were analyzed using High Sensitivity DNA chips (Agilent), a 10 nM normalized library was pooled, and sequencing performed on a NovaSeq S2 (Illumina) with 28 and 75 bases for reads 1 and 2 and 10 bases each for index reads 1 and 2.

snRNA-seq libraries were prepared from 6 WT and 6 *Srrm2*^+/-^ mice for each brain region.

##### RNAScope and QuPath Quantification

Brains from 6 WT and *Srrm2*^+/-^ male P29 littermates were sectioned sagittally at 10 μm thickness. One slide contained three sections from one animal. RNAscope™ Multiplex Fluorescent V2 Assay was conducted following the manufacturer’s protocol for one slide per animal. The probe and fluorophore configuration was C1-*Aspa-*Opal 680, C2-*Plp1*-Opal 570, C3-*Ppp1r1b*-Opal 520. Slides were imaged using the ZEISS Axioscan 7 and Zen 3.7 imaging software. Quantification was performed using QuPath v0.6.0 software. An oval ROI of 1.2×10^6^ um^2^ was randomly placed in the dorsal area of the striatum. For the PFC, a hand-drawn ROI based on the Allen Brain Atlas was created following the curvature of the cortex adjacent to the olfactory bulb and the corpus callosum (see Fig S4). Cell detection was performed using QuPath’s built-in cell detection feature with pixel width/height = 0.32 μm based on image dimensions. Trained classifiers using QuPath’s “Random trees (RTrees)” method on mean intensity in the nuclei and cytoplasm were used to classify cells as positive or negative for *Aspa, Plp1,* or *Ppp1r1b.* The classifier was trained using an equal number of ∼8-10 cells per condition (e.g. *Aspa^+^, Aspa^-^)* on one section per animal. A composite classifier was created to collect data on collocalization (e.g. *Aspa^+^Plp^+^Ppp1r1b^-^)*. ROI placement and classifier training was done blinded to genotype.

##### EEG implantation and recording

EEG electrodes were implanted as described previously ^45,49^. Briefly, 8- to 10-week-old mice (n = 13 WT and 17 *Srrm2*^+/-^) were deeply anesthetized with isoflurane. A prefabricated EEG/EMG headmount (#8201SS, Pinnacle Technology, Lawrence, KS) was secured to the skull with four 0.10’’ intracranial electrode screws (#8403, Pinnacle Technology) at the following stereotactic coordinates: parietal recording electrode (-1.5 AP, 2.0 ML to Bregma), frontal recording electrode (1.5 AP, 1.5 ML to Bregma), and ground and reference electrodes (bilaterally -1 AP, 2 ML to Lambda). The electromyogram (EMG) electrodes were placed bilaterally in the nuchal muscles. Electrodes were soldered to the EEG/EMG head-mount and dental acrylic was used to secure the connections. Animals were given at least two weeks of post-operative recovery before EEG recording. Following recovery from EEG implantation, mice were tethered to the Pinnacle recording system, with at least 24 hours of habituation before recording. For sleep/wake recordings, EEG/EMG signals were recorded for 24 hours from the onset of the dark phase, ZT12 (6 pm EDT or 7 pm EST). Animals remained tethered to the Pinnacle system throughout the testing period with *ad libitum* access to food and water. All signals were digitized at a sampling rate of 1000 Hz, filtered (1–100 Hz bandpass for EEG; 10–1 kHz bandpass for EMG), and acquired using the Sirenia Acquisition program (Pinnacle Technology). EEG recordings were carried out at ∼3- and ∼6-months of age.

##### Behavior profiling

Investigators were blinded to the genotypes for all behavioral tests.

##### Homecage locomotor activity monitoring

Mice were transferred to Digital Ventilated Cages (DVC, Tecniplast) and were single housed throughout the duration of the experiment. Briefly, DVC racks use 12 electrodes beneath each cage that radiates an electromagnetic field that can detect disturbances that correlate with animal movement. The DVC Analytics web platform was used to generate animal locomotor indices, which calculate the percentage of 12 electrodes that measured movement over a 1-minute bin. For the locomotor activity index, a value of 1 means 12 electrodes measured movement, and a value greater than 1 means that there were more than 12 detected movements. A running wheel was added to the cage in the final week of the experiment.

##### Open field

Mice were monitored using the SuperFlex Open Field system (40 cm x 40 cm x 40 cm; Omnitech Electronics, Inc., Columbus, OH) for 60 minutes. The animals’ position was captured in real time using Fusion system software (Omnitech Electronics, Inc.). Total distance traveled, center/margin distance, and jump counts were calculated using the system’s inbuilt software. The center was defined as the central 8 x 8 squares (of a 16 × 16 matrix) plus the coordinates between the outermost beams of the area and the adjacent non-area beams, whereas the margin was defined as the exterior 4 x 16 and 16 x 4 matrices for the left, right bottom and top regions of the arena.

##### Forced swim test

Mice were placed gently by their tail in a 4-liter Griffin beaker filled two-thirds with water maintained at room-temperature (24°C ± 1°C) for 6 minutes, and the amount of time the mice spent immobile was recorded. The temperature was tested before each test with an infra-red or glass thermometer.

##### Acoustic startle and PPI

To test acoustic startle, the test mouse was placed in a sound-attenuating chamber on a motion-sensitive platform. The background noise was set at 60 dB. After a 5-minute period of acclimation, a series of sudden, loud acoustic stimuli of 60, 70, 80, 85, 90, 95, 100, 105, 115 dB were presented in random order. For PPI, mice were exposed to 20 millisecond-long pre-pulses of 74, 78, 82, 86, 90 or dB. 100 milliseconds after the pre-pulse, mice were exposed to a constant startle tone of 105 dB that lasted 40 milliseconds. For both tests, the intertrial interval, which was randomly chosen, was 10, 15, 20, 25, or 30 seconds. For each stimulus, 5 trials were performed (in random order, with random order of inter-trial interval). The startle response was recorded as the force exerted by the mouse in response to each acoustic stimulus. The recording time was set to 250 milliseconds. The percentage of prepulse inhibition of the startle response was calculated as 100 × [startle response_(pre-pulse)_ – (startle response_(prepulse + pulse)_]/startle response_(pre-pulse)_.

##### Novel objection recognition

Novel object recognition testing was performed in a 30 cm (w) X 30 (l) X 10 (h) arena made of acrylic. To reduce anxiety in the animals, light in the testing area was dimmed to 30 ± 5 lux. The testing was conducted over two days. On day 1 mice were habituated to the empty arena for 15 minutes. On day 2 mice were exposed to two identical objects in the rear left and right corners of the arena for 10 minutes. The mice were then returned to their home cages. After 10 minutes or 24 hours, mice were returned to the test arena with one of the familiar objects in place and a novel object of the same size in the same area as the other familiar object. The amount of time the mice spent interacting with the novel and familiar objects was recorded; testing was carried out for a period of 5 minutes. The arena and objects were cleaned between trials to prevent odor cues. Animal tracking and object exploration scoring was performed with Ethovision XT17 (Noldus).

##### Social interaction and social novelty

Social interaction and preference for novel animals were evaluated using the standard 3-chambered apparatus, which consists of a central chamber connected to two side chambers. Mice were placed in the central chamber and allowed to explore all three chambers freely for five minutes to habituate to the arena. After the habituation period, an unfamiliar male mouse that had no prior contact with the test mouse, was placed in one of the side chambers enclosed in a small round wire cage. An identical empty wire cage (the “object”) was placed in the other chamber. The animals serving as strangers were male mice that had previously been habituated to placement in the small cage. Both doors to the side chambers were then unblocked and the test mouse was allowed to explore the entire arena for a 10-minute session. The amount of time spent investigating the stranger mouse by the test mouse was recorded manually using a stopwatch. Social interaction was quantified by (time spent interacting with the mouse) / (time spent interacting with mouse + time spent interacting with object). After a 10-minute trial, a second stranger mouse was placed in the empty wire cage. The amount of time spent investigating each of the mice by the test mouse was recorded manually, and preference for novel animals was calculated as (novel mouse time) / (novel mouse time + familiar mouse time).

##### Y-maze

Forced alternation tests were conducted using a symmetrical Y-maze, which consists of a starting arm, an open arm, and a novel/blocked arm. Each arm of the Y-maze was 35 cm long, 5 cm wide and 10 cm high. Mice were habituated to the testing room for at least 30 min. Following habituation, mice were placed in the starting arm and allowed to explore two arms (starting and open) of the Y-maze for 10 minutes, while the entrance to the third arm was blocked. After the sample trial, the mouse was returned to its home cage for a 30 min inter-trial interval. The block in arm 3 was then removed and the mouse was placed into the starting arm again and allowed access to all three arms of the maze for five minutes. Mice were required to enter an arm with all four paws for it to be counted as an entry. The time spent in the novel arm was calculated as a percentage of the total time in novel and open arms during the 5-min retrieval trial. The maze was cleaned between trials.

##### Grip strength

Forelimb grip strength was assessed using a weighted-bag apparatus connected to a wire mesh. Mice were held by the base of the tail and allowed to grasp the mesh with their forepaws. Once a firm grip was established, the mouse was lifted vertically until the weights cleared the bench. A successful trial was defined as a sustained hold of least 3 seconds.

Testing began with a baseline weight of 10 g, followed by 10 g increments for each successful lift. Mice were granted three trials per weight level, with a 20-second inter-trial rest period. The maximum weight successfully lifted for 3 seconds was recorded as the final score.

### QUANTIFICATION AND STATISTICAL ANALYSIS

#### Bulk RNA-seq gene DE

*RSEM* v1.3.084 ^74^ was used to estimate gene and isoform expression values. The M25 (GRCm38.p6) GENCODE reference was generated using RSEM-prepare-reference with default parameters. Expression values were calculated using RSEM-calculate-expression with the following flags: –bowtie2, –paired-end, –estimate-rspd, –append-names, and –sort-bamby-coordinate. *DESeq2* v1.20.0 ^75^ was used to perform DE analysis for each brain region and age. For comparisons of normalized counts across regions and age, all samples of every region and age were read into a single DESeq object. Log2FC values were adjusted using DESeq2’s *lfcShrink* function with the ‘normal’ shrinkage estimator.

#### Age-dependent compensation analysis

To identify genes exhibiting age-dependent compensation, an interaction analysis was performed using a generalized linear model within the DESeq2 framework. A comprehensive design formula, *∼ Region + Age + Genotype + Age:Genotype*, was used. In this model, *Region* was included as a covariate to account for baseline transcriptomic differences across the eight brain regions. The *Age:Genotype* interaction term was specifically employed to identify genes where the transcriptomic difference between *Srrm2*^+/-^ and wild-type mice was significantly altered between the 1-month and 3-month timepoints. GSEA was performed on the resulting ranked list of genes to identify biological pathways undergoing significant temporal shifts across regions in *Srrm2*^+/-^ mice brains.

#### Bulk RNA-seq isoform DE and P-value aggregation

Reads were pseudoaligned to either Gencode vM32 (mouse) or v47 (human) transcriptomes using *Kallisto* 0.46.1 ^76^, with the number of bootstraps set to 50 (mouse) or 10 (human). Isoform-level DE analysis was performed using *Sleuth* 0.30.1 ^77^ by comparing a full model (with genotype as the only parameter) with a reduced model for each brain region and age. *P*-values generated by the isoform DE tests for each gene were aggregated (meta-analyzed) using the Lancaster method, as described in Yi et al. ^32^, also using *Sleuth*. Gene-level mRNA abundance estimates do not fully capture the dynamics of its constituent isoforms. For example, the abundance of different isoforms of the same gene changing in opposite directions can cancel each other out and lead to minimal changes in overall mRNA abundance (the sum of the abundances of constituent isoforms). In such a case, the aggregation of the *P*-values of each constituent isoform—which is performed by this method—results in an even lower *P*-value thereby unmasking and prioritizing genes with dynamic effects among its multiple constituent transcripts ^32^.

#### Leafcutter and rMATS splicing analyses

RNA-seq FASTQ files were aligned to the mm39 mouse genome (or hg38 for human iPSC-derived neurons) using STAR in a comprehensive two-pass mode. Briefly, an initial alignment was performed to identify splice junctions; after filtering for annotated, non-canonical, and low-support junctions (≤ 2 reads), the remaining junctions were provided via the *--sjdbFileChrStartEnd* flag for a second, high-sensitivity alignment pass.

To quantify differential intron excision, exon-exon junctions were extracted from the resulting BAM files using regTools and analyzed via Leafcutter ^79^. Intron clustering and differential excision analysis were performed as previously described, with altered splice junctions visualized using *leafviz* ^79^ and *ggsashimi* ^80^.

Complementary differential alternative splicing analysis was performed using rMATS-turbo (v4.3.0) ^81^. The analysis was conducted in paired-end mode (*-t paired*) with a read length of 50 bp and the *--allow-clipping* flag enabled. rMATS-turbo was used to quantify five primary types of alternative splicing: skipped exons (SE), alternative 5’ splice sites (A5SS), alternative 3’ splice sites (A3SS), mutually exclusive exons (MXE), and retained introns (RI). For all analyses, differential splicing events were considered significant at a False Discovery Rate (FDR) <= 0.05. Differential junction counts and exon counts (“JCEC”) were analyzed using the summary output file.

#### High-resolution isoform analysis of Syngap1

*Syngap1* mRNA isoforms (Gencode M25) were assigned to the four UniProt SynGAP protein isoforms based on shared C-terminal splice architecture. Mapping was guided by Gou et al. ^31^ This approach was necessary due to the structural complexity of the *Syngap1* locus, which generates diversity through both alternative transcription start sites (N-terminal) and alternative C-terminal splicing. Because transcript-level quantification distributes reads across shared exonic regions among isoforms with overlapping sequence, individual transcript counts may underestimate the total functional output of a given splice variant. Accordingly, transcripts encoding previously defined C-terminal isoforms were summed to generate transcriptional pools corresponding to C-terminal isoforms detected by proteomics.

#### *Syngap1* mapping

Gamma (A0A0A6YVS6: *Syngap1* 203, 210); Alpha1 (A0A0J9YUM2: *Syngap1* 202, 208, 212, 213); Alpha2 (F6SEU4: *Syngap1* 201, 204, 214); Beta (A0A0J9YVH8: *Syngap1* 207). *Agap3* mapping: Agap3-FL (Q8VHH5: *Agap3* 201, 202); Crag (A0A1D5RMG4: *Agap3* 203–206).

This aggregation reconciles the greater number of annotated Gencode transcript isoforms with the four SynGAP protein isoforms represented in the UniProt database used for proteomic analysis. *Syngap1* 210, which was not included in Gou et al. (likely due to differences in annotation versions), was classified as Gamma because manual inspection confirmed the presence of the in-frame exon specific to the Gamma C-terminus.

For cortex proteomic–transcriptomic effect size comparisons, PFC and SSC RNA-seq samples were pooled to increase statistical power. Transcriptome-wide isoform-level differential expression analysis was then performed using DESeq2 in R. To account for baseline expression differences between PFC and SSC while isolating the effect of genotype, a multi-factor design (∼ Region + Genotype) was employed. Log2 fold changes (*Srrm2*^+/-^ vs. WT) from isoform-level RNA differential expression were correlated with corresponding proteomic log2 fold changes. Analyses included age-matched 1-month RNA and proteomics datasets, as well as 3-month RNA compared to 4-month proteomics (denoted “3/4mo”). Isoform-level RNA log2 fold changes used for this analysis were unshrunken to preserve effect size magnitude.

#### Single-nucleus RNA-seq analysis

For striatum samples, the 10X Genomics *Cell Ranger* v7.2.0 ^93^ pipeline was used to align reads from snRNA-seq to a version of the mouse mm10 reference transcriptome that includes introns (refdata-gex-mm10-2020-A, 10X Genomics). The –chemistry=SC3Pv3 and –include-introns=TRUE flags were used in addition to default parameters for Cell Ranger count. For PFC samples, Cell Ranger v3.0.2 was used with the same reference transcriptome. Ambient RNA removal was performed on striatum samples using CellBender v0.3.0 ^82^. Unique Molecular Identifier (UMI) counts were analyzed using the *Seurat* v4.4.0 package ^94^. Nuclei expressing less than 500 genes were removed and the remaining nuclei were normalized by total expression, multiplied by ten thousand, and log transformed. Seurat’s *ScaleData* function was then used to scale the data. Linear dimension reduction by PCA was performed using Seurat’s RunPCA function on the scaled data on variable genes, and significant PCs were identified. The nuclei were clustered in the PCA space using Seurat’s FindNeighbors and FindClusters functions then visualized using Uniform Manifold Approximation and Projection (UMAP) using significant PCs. Doublet identification was performed using the *Scrublet* v0.2.3 Python package^84^ with default parameters. Clusters and nuclei with a scrublet score above the region-specific thresholds (0.40 for 1-month PFC and 0.41 for 1-month striatum) were determined to be doublets and were removed from the data.

UMAP clusters were assigned to cell type by manual examination of enrichment/depletion of brain-region-specific cell type marker genes. Neuronal subtypes were identified by re-clustering the nuclei of a major cell type and evaluating expression of subtype-specific marker genes. Marker genes were obtained from dropviz.org ^24^.

#### Cell composition analysis

Cell composition analysis was performed in R using the *speckle* v1.0.0 ^87^. The *propeller* function with *arcsin* normalization was used to test for statistical differences in major cell type and subtype composition between genotypes in each examined brain region. For striatum samples, cell composition analysis was performed using either inhibitory neurons as a whole or after dividing inhibitory neurons into subtypes (dSPN, iSPN, eSPN, PV interneurons).

#### Single-nucleus RNA-seq DE

DE analysis (*Srrm2*^+/-^ vs. WT) was performed using the scOnline pseudocell method ^85,86^ for each cell type and subtype in each brain region. By aggregating the expression of a group of nuclei with similar transcriptomes within each replicate, cell type, and subtype, this method resolves known technical variation issues of snRNA-seq transcriptome data including low capture rates from dropouts and pseudoreplication, while avoiding the pitfalls of pseudobulk approaches, such as low statistical power and high variation in sample sizes ^85,86^. To construct pseudocells, single nuclei were grouped by cell type for each sample. Within each group, pseudocell centers were identified by applying k-means clustering so that each cluster contained 20 nuclei at minimum and 40 nuclei on average. Cell types and subtypes that were identified but did not have enough nuclei present to create at least two pseudocells per sample were removed. To generate the pseudocell counts matrix, we aggregated the raw UMI counts of cells assigned to each pseudocell. Differential expression analysis was performed using limma’s *duplicateCorrelation* mixed model analysis function with robust empirical Bayes moderated t-statistics and mouse ID as random effect. Genotype and the percentage of mitochondrial reads were used as covariates for all analyses. Genes with FDR-adjusted *P*-value of < 0.01 were defined as DE genes.

#### Proteomics data analysis

Mass spectra were analyzed using Spectrum Mill MS Proteomics Software (Broad Institute) with a mouse database from Uniprot.org downloaded on 04/07/2021 containing 55734 entries. Search parameters included: ESI Q Exactive HCD v4-35-20 scoring, parent and fragment mass tolerance of 20 ppm, 40% minimum matched peak intensity, trypsin allow P enzyme specificity with up to four missed cleavages, and calculate reversed database scores enabled. Fixed modifications were carbamidomethylation at cysteine. TMT labeling was required at lysine, but peptide N termini could be labeled or unlabeled. Allowed variable modifications were protein N-terminal acetylation, oxidized methionine, pyroglutamic acid, and pyro carbamidomethyl cysteine. Protein quantification was achieved by taking the ratio of TMT reporter ions for each sample over the TMT reporter ion for the median of all channels. TMT18 reporter ion intensities were corrected for isotopic impurities in the Spectrum Mill protein/peptide summary module using the afRICA correction method which implements determinant calculations according to Cramer’s Rule and correction factors obtained from the reagent manufacturer’s certificate of analysis (https://www.thermofisher.com/order/catalog/product/90406) for lot numbers VH310017 and WG333575. After performing median normalization, a moderated two-sample t-test was applied to the datasets to compare WT and *Srrm2*^+/-^ sample groups at 1- and 4-months of age.

#### Gene set enrichment analysis

GSEA was performed with the R Bioconductor package *fgsea* ^88^. Genes/proteins were pre-ranked using the appropriate test-statistics (DESeq2 Wald test-statistic for bulk RNA-seq, and the moderated t-test statistic for single-nucleus RNA-seq and synapse proteomics experiments). We carried out GSEA using two independent gene set collections—M5 ontology gene sets (v2023.2) obtained from the Molecular signatures database (MSigDB) ^19^, and (ii) SynGO gene sets (v1.2) ^20^ combined with other curated gene sets, which include SCZ GWAS hits ^22^, SFARI ASD risk genes ^95^, bipolar GWAS hits ^21^. Also combined with SynGO sets were the ‘activity regulated’ gene-set, which consisted of neuronal activity-induced rapid, delayed and slow primary response genes defined by Tyssowski et al. ^96^, and the ‘dopamine-induced’ gene-set, consisting of genes with FDR-adjusted *P*-value < 0.1 that were upregulated in striatal neuronal culture following one hour dopamine treatment ^97^. For bulk RNA-seq we also specifically assessed the “LEIN_OLIGODENDROCYTE_MARKERS” set from mSigDb. For proteins with multiple isoforms, the isoform with the largest effect-size was used for pre-ranking. Because the enrichment analyses were carried out using the proteins’ gene symbols, we use the terms gene set and protein set interchangeably in this study.

#### EEG data analysis

All raw recordings were segmented into 10-second epochs, and each epoch was classified into NREM, REM, or wake stages using the Light Gradient Boosting Machine (LightGBM) algorithm, as previously described ^46^. Sleep spindle detection was performed using the Luna toolbox, which uses a wavelet-based detection method ^98^. Spindle density, amplitude, integrated spindle activity (ISA), and spindle duration were calculated for all animals based on the detection threshold of WT animals. Spindle parameters and power density data were then analyzed using a linear mixed-effects model (ex. *spindle density ∼ genotype* frequency + subjectID*) performed in Python. The statistical impact of genotype on measures was based on likelihood ratio test between two models: (*spindle density ∼ genotype* frequency + subjectID*) and (*spindle density ∼ frequency + subjectID*). Prior to statistical analysis, spindle density outliers were removed using stringent 2IQR (interquartile range method), where values below Q1 - 2IQR or above Q3 + 2IQR were considered outliers. For each subject and genotype group (WT, *Srrm2*^+/-^), spindle parameters and EEG power were averaged across conditions and grouped by subject ID, genotype, and frequency band (F). Spindle detection was performed using different center frequencies (9, 11, 13, 15 Hz). For EEG power analysis, we examined different frequency bands, including slow oscillations, delta, theta, alpha, sigma, beta, and gamma bands. The statistical significance of differences across genotypes and frequency bands was evaluated using *P*-values from the mixed model output, with a significance threshold set at *P* < 0.05.

### ADDITIONAL RESOURCES

No additional resources are reported.

